# Mutant FGFR3 restricts bone yet expands cortex via ERK-mediated self-repression

**DOI:** 10.64898/2026.02.03.703675

**Authors:** Zhuangzhi Zhang, Zhejun Xu, Tongye Fu, Wenhui Zheng, Zizhuo Sha, Chuannan Yang, Feihong Yang, Jialin Li, Jing Ding, Zhengang Yang

## Abstract

*FGFR3* gain-of-function mutations cause achondroplasia ^1,2^, the most common form of dwarfism, yet trigger paradoxical cerebral overgrowth and skeletal stunting ^3–9^. Here, we demonstrate that *Fgfr3-K644E* mutant drives enhanced ERK activation, which then represses *Fgfr3* expression and downregulates ERK signaling. In the developing cortex, this mechanism transiently expands cortical stem cells by activating ERK, then shifts signaling to downregulate *Fgfr3*/ERK/PKA and enhance YAP/TAZ, promoting cerebral overgrowth and premature ependymal maturation. In growing bones, *Fgfr3-K644E* mutant first elevates ERK to inhibit chondrocyte pre-hypertrophy, then downregulates *Fgfr3/*ERK/PKA and upregulates YAP/TAZ to accelerate premature hypertrophy and ossification. Our findings establish that *FGFR3* mutations limit bone yet expand cortical size via an ERK negative feedback loop, revealing a unified disease mechanism and novel therapeutic targets.

## Main Text

The discovery three decades ago that *FGFR3* gain-of-function (GOF) mutations cause achondroplasia—the most common skeletal dysplasia—opened a new era in understanding human growth disorders ^1,2^. Since then, mutations in the same receptor tyrosine kinase have been linked to a spectrum of related skeletal conditions, including hypochondroplasia, SADDAN, and thanatophoric dysplasia (TD), all characterized by impaired cartilage development. While *FGFR3* GOF mutations potently inhibit longitudinal bone growth, they conversely promote brain overgrowth, as notably observed in individuals with TD ^3–9^. This tissue-specific duality reveals a key gap: how do identical FGFR3 mutations yield opposite skeletal versus cerebral effects? The molecular basis for this context-dependent signaling remains unknown, hampering targeted therapies. Here, we address this paradox by mapping FGFR3 effector regulation across developmental contexts.

## Results

### The *Fgfr3-K644E* GOF mutation suppresses *Fgfr3* expression

The *Fgfr3-K644E* GOF mouse model (*EIIa-Cre; Fgfr3^K644E/+^*) established in this study carries a *K644E* knock-in mutation, which corresponds to the human *K650E* mutation found in TDII (Extended Data Fig. 1). This model reveals a critical role for *Fgfr3*, which phenotypically drives posterior cortical expansion and results in severe, lethal skeletal dysplasia at birth ^6,8,10–13^ (Extended Data Fig. 1).

To investigate the molecular mechanism underlying cortical expansion in *Fgfr3-K644E* mice, we performed scRNA-Seq on E12.5 cortices (Fig. 1a-c). Gene expression analysis in cortical radial glia (RGs) revealed that ERK signaling downstream targets, including *Bmp7*, *Etv1/4/5*, *Spry2*, and *Hopx,* were upregulated in *Fgfr3-K644E* mice compared with controls (Fig. 1d). In contrast, genes associated with YAP signaling (YAP/TAZ activity) (*Amotl2, Cyr61, Wwc2, Wwtr1, Yap1*) and SHH signaling (*Boc, Cdon, Gas1, Gli2, Gli3*) were downregulated (Fig. 1d). Moreover, the downregulation of *Nr2f1* and *Fgfr3* suggests that the *Fgfr3-K644E* mutation suppresses its own expression through enhanced ERK signaling (Fig. 1d) ^14^. Consistent with ERK-mediated repression of WNT signaling ^15^, *Lef1* and *Axin2* in RGs were also reduced (Fig. 1d). Moreover, enhanced self-renewal of cortical RGs by ERK was accompanied by downregulation of neuronal progenitor genes such as *Eomes* and *Neurog2* (Fig. 1d). HOPX is a well-established downstream target gene of ERK in cortical RGs ^14^. Immunostaining and in situ hybridization of the E12.5 cortex revealed that caudal RGs displayed a slight increase in pERK, accompanied by strong HOPX expression and reduced *Fgfr3* expression (Fig. 1e). These observations were further validated by Western blot analysis (Fig. 1f, g) ^10^.

**Fig. 1.**
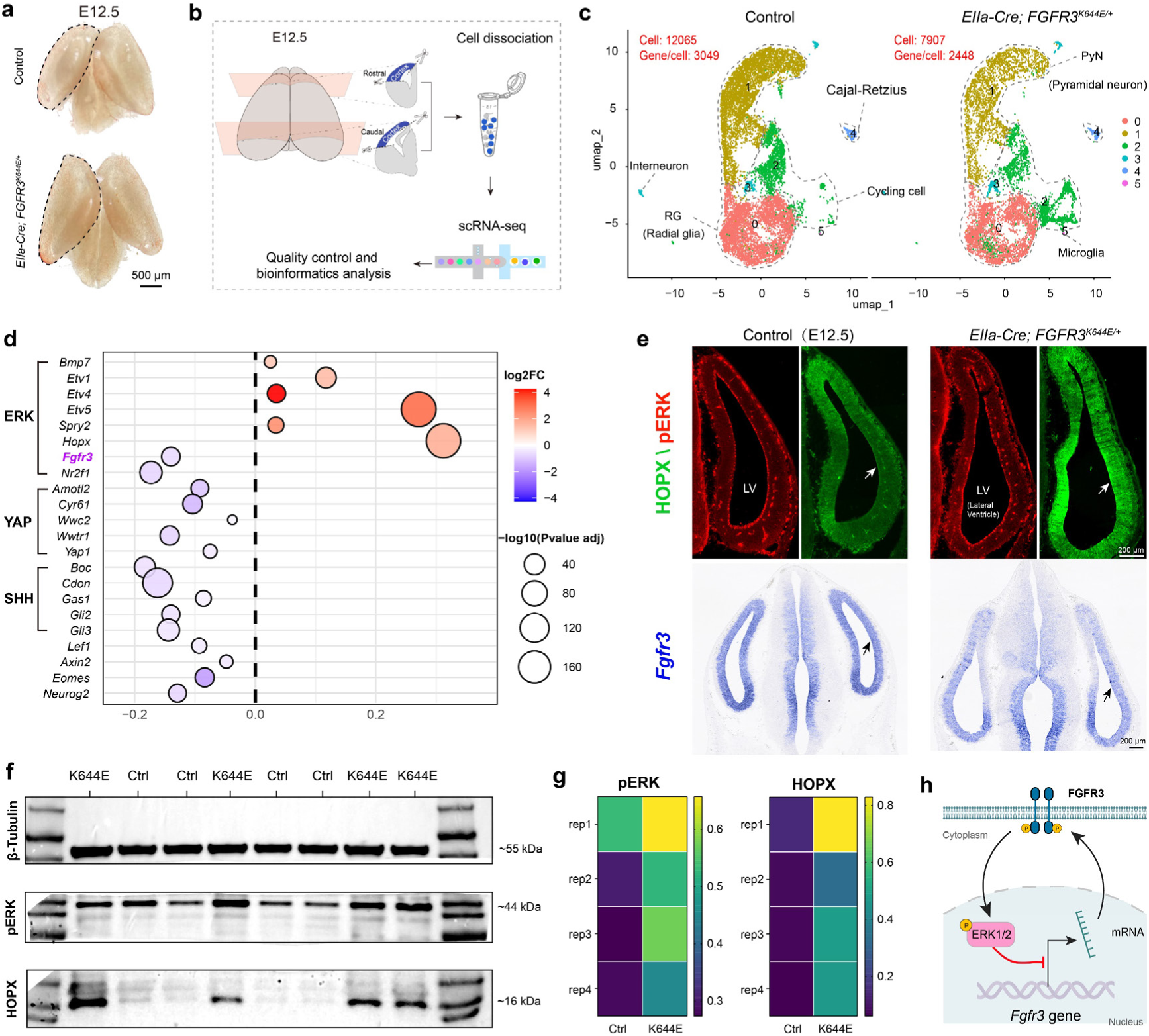
The *Fgfr3-K644E* mutation initiates a negative feedback loop that suppresses its own transcription via ERK signaling. **a-c,** scRNA-Seq analysis of cortical RGs (stem cells) in control and *Fgfr3-K644E* mice at E12.5. **d,** ERK signaling genes (*Bmp7, Etv1/4/5, Spry2, Hopx*) were upregulated in *Fgfr3-K644E* cortical RGs, while *Fgfr3*, *Nr2f1*, YAP/TAZ- and SHH pathway-related genes were downregulated. **e,** At E12.5, immunostaining and in situ hybridization revealed a subtle increase in pERK alongside strong HOPX expression and a downregulation of *Fgfr3* in RGs of the caudal cortex (arrows). **f, g,** Western blotting confirms elevated pERK and HOPX in dissociated cortical tissue from *Fgfr3-K644E* mice at E12.0. **h,** The *Fgfr3-K644E* mutation first triggers ERK signaling, which then gives rise to *Fgfr3* transcriptional self-suppression, a process that subsequently mediates downregulation of the ERK pathway.

We next investigated the functions of the *FGFR3-K644E* mutation and constitutive ERK signaling (*MEK1DD* allele) using conditional approaches. Overall, at E12.5, both *Emx1-Cre; Fgfr3^K644E/+^*, *Emx1-Cre; Fgfr3^K644E/K644E^*, and *Emx1-Cre; Rosa^MEK1DD/+^*mice exhibited phenotypes similar to those of *Fgfr3-K644E* mice, including cerebral cortex enlargement and pronounced upregulation of HOPX expression (Extended Data Fig. 2a, d). While *Emx1-Cre; Rosa^MEK1DD/+^* mice exhibited strong and sustained pERK expression, pERK levels were only transiently elevated in cortical RGs of *Emx1-Cre; Fgfr3^K644E/K644E^* mice (Extended Data Fig. 2a-d). *In situ* hybridization analysis further revealed that E12.5 cortical RGs in *Emx1-Cre; Fgfr3^K644E/K644E^* mice exhibit upregulated *Spry2* alongside downregulated *Fgfr3* and *Nr2f1* (Extended Data Fig. 3a-d). Taken together, these results demonstrate that the *Fgfr3-K644E* mutation induces a transient enhancement of ERK signaling in cortical stem cells, as activated ERK represses *Fgfr3* expression ^14^, thereby limiting its own activation through a negative feedback loop (Fig. 1h).

### *Fgfr3-K644E* downregulates ERK signaling in perinatal cortical RGs

To further assess the long-term impact of the *Fgfr3-K644E* mutation, we performed scRNA-seq on E15.5 cortices (Extended Data Fig. 4a, b). While cortical expansion was more pronounced in mutants compared to controls and relative to E12.5, no significant changes were detected in PKA-, SHH-, or YAP-related genes, and ERK signaling upregulation in cortical RGs was no longer evident (Extended Data Fig. 4c).

scRNA-Seq of E18.5 *Fgfr3-K644E* cortices revealed that most ERK downstream targets remained unchanged in RGs (Extended Data Fig. 5a-d). However, *Hopx* was markedly downregulated (Extended Data Fig. 5d), indicating downregulation of ERK signaling. Although ERK and PKA signaling can mutually reinforce each other ^16–19^, at E18.5, no changes were detected in the expression of PKA-related genes (*Adcyap1r1, Dio2*) (Extended Data Fig. 5d). In contrast, components of the SHH pathway (*Cdon, Gas1*) and its downstream targets in cortical RGs (*Ascl1, Olig1, Olig2*) were downregulated (Extended Data Fig. 5d). The most pronounced change was the upregulation of markers reflecting YAP/TAZ activity (*Amotl2, Cyr61, Ctgf*) (Extended Data Fig. 5d). Previous studies have established that YAP signaling is essential for ependymal cell formation ^18,20,21^. Consistent with this, we observed elevated expression of ependymal-associated genes (*Anxa2, Enkur, Gja1, Rfx3, S100a6, Sox9*) in cortical RGs (Extended Data Fig. 5d). Immunostaining further confirmed a marked downregulation of HOPX along with a pronounced upregulation of SOX9 expression in E18.5 cortical RGs (Extended Data Fig. 5e, f). Notably, SOX9 is known to promote ependymal cell maturation ^22^.

Given the neonatal lethality of *Fgfr3-K644E* mice, we performed scRNA-Seq analysis on P2 *Emx1-Cre; Fgfr3^K644E/K644E^*homozygous mice. This revealed a pronounced downregulation of ERK, PKA, and SHH signaling in cortical RGs (Fig. 2a-f). Conversely, YAP signaling and early ependymal cell genes were markedly upregulated (Fig. 2f). This transcriptional shift correlated with an increase in ependymal cell generation by P2 (Fig. 2e). Immunostaining further demonstrated enhanced nuclear localization of YAP/TAZ, accompanied by a greater number of FOXJ1-positive ependymal cells lining the cortical ventricle (Fig. 2g, h). Thus, ERK signaling is reduced in perinatal cortical RGs *of Fgfr3-K644E* mutant mice, leading to attenuated PKA and enhanced YAP/TAZ activation and premature ependymal differentiation.

**Fig. 2.**
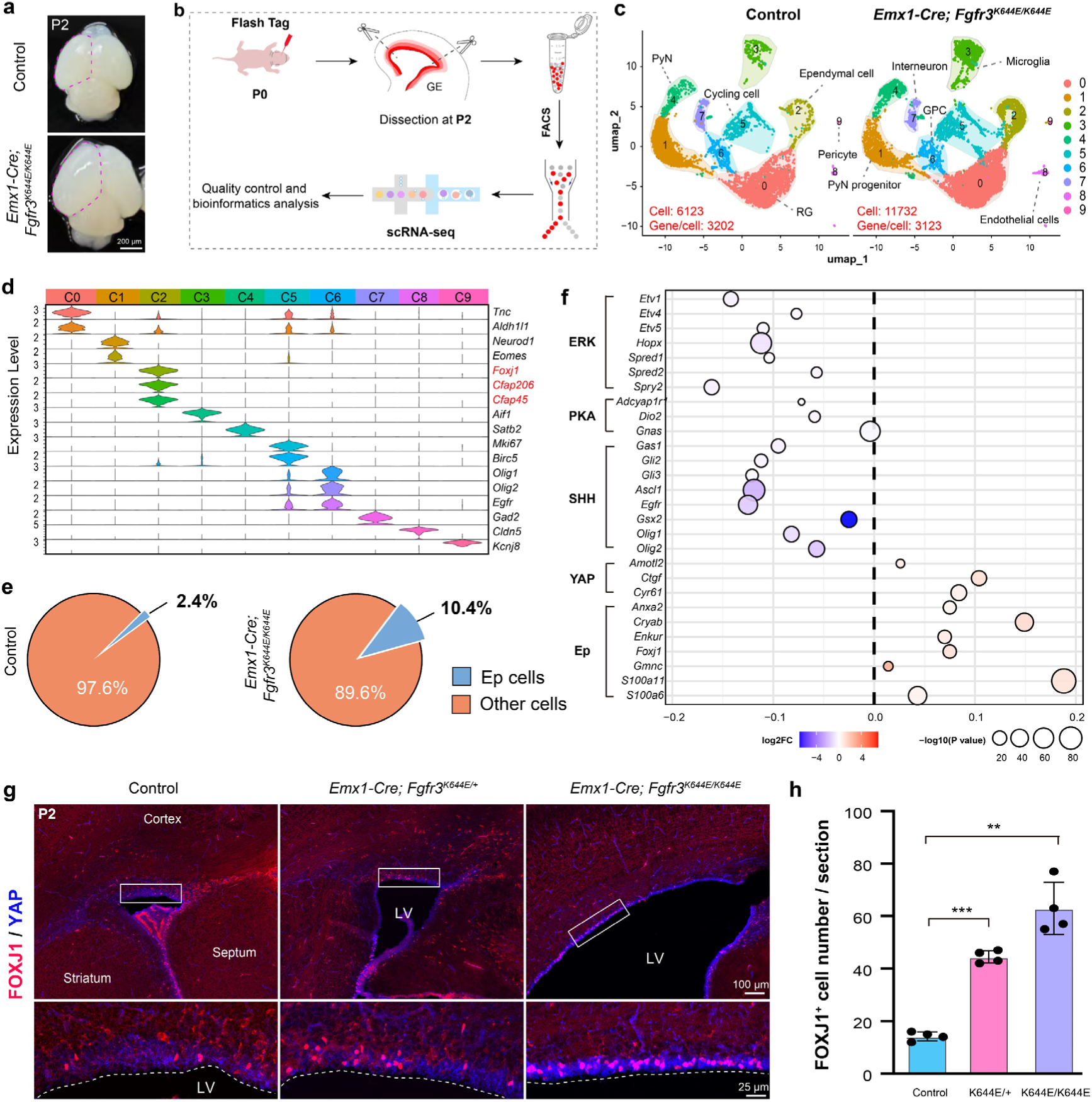
The *Fgfr3-K644E* mutation downregulates ERK signaling in P2 cortical RGs. **a-c,** scRNA-Seq analysis of cortical RGs in control and *Emx1-Cre; Fgfr3^K644E/K644E^*mice at P2. **d,** Violin plots show the expression levels of marker genes. **e,** The ependymal (Ep) cell population was expanded. **f,** Genes associated with ERK, PKA, and SHH-SMO signaling were downregulated in mutant cortical RGs (cluster 0), whereas genes in the YAP signaling and ependymal cell markers were upregulated. **g, h,** Immunostaining showed increased nuclear YAP/TAZ localization and more FOXJ1+ cortical ependymal cells. n = 4; mean ± SEM; **, P<0.01; ***, P<0.001 (one-way ANOVA with Tukey’s post-hoc test).

### *Fgfr3-K644E* mutation and constitutive ERK signaling drive differences in long-term survival rates

*Fgfr3-K644E* mutation and constitutive ERK signaling each enlarge the cortex, but in *Emx1-Cre; Fgfr3^K644E/+^* mice, ependymal cells developed essentially normally and the animals survived long-term (Extended Data Fig. 6a-c). Conversely, *Emx1-Cre; Rosa^MEK1DD/+^* mice failed to generate differentiated cortical ependymal cells and typically died within three months (Extended Data Fig. 6d, e). Similarly, *Nestin-Cre; Fgfr3^K644E/+^* mice displayed well-developed ependymal cells and survived long-term (Extended Data Fig. 7a-d). In contrast, *Nes*-*Cre; Rosa^MEK1DD/+^*mice, with persistently elevated ERK signaling in forebrain neural stem cells, displayed defective ependymal development that led to severe hydrocephalus and death within three weeks (Extended Data Fig. 7e-h).

In summary, unlike constitutive MEK1DD activation, *Fgfr3-K644E* mutation triggers only transient early ERK activation in cortical RGs, which is then downregulated through a self-limiting negative feedback loop that suppresses *Fgfr3* expression (Fig. 1h). The consequent attenuation of PKA and amplification of YAP signaling collectively drive the observed phenotypic outcomes. This fundamental understanding provides crucial insights for elucidating the mechanisms by which *Fgfr3* mutations influence bone development.

### *Fgfr3* expression mirrors YAP/TAZ signaling in chondrocytes

While *FGFR3* GOF mutations are responsible for dwarfism, the exact molecular mechanisms are still not completely understood. ^3–5,8,9^. To address this, we performed scRNA-Seq on developing humeri collected from control and *Fgfr3-K644E* mice across 6 time points, from E13.5 to E18.5 (Fig. 3a). A total of 12 samples were analyzed, yielding 116,523 cells with an average of 2,751 genes detected per cell (Fig. 3b). This effort established a comprehensive single-cell transcriptomic atlas of humerus development and a dedicated database for the *Fgfr3-K644E* mutant model (Fig. 3b, c). Next, we isolated the chondrocyte lineage for in-depth analysis, encompassing proliferating chondrocytes (PCs, including both round and flat PCs), prehypertrophic chondrocytes (PHCs), hypertrophic chondrocytes (HCs), as well as osteoblasts derived from both HCs and mesenchymal stem cells (Fig. 3d).

**Fig. 3.**
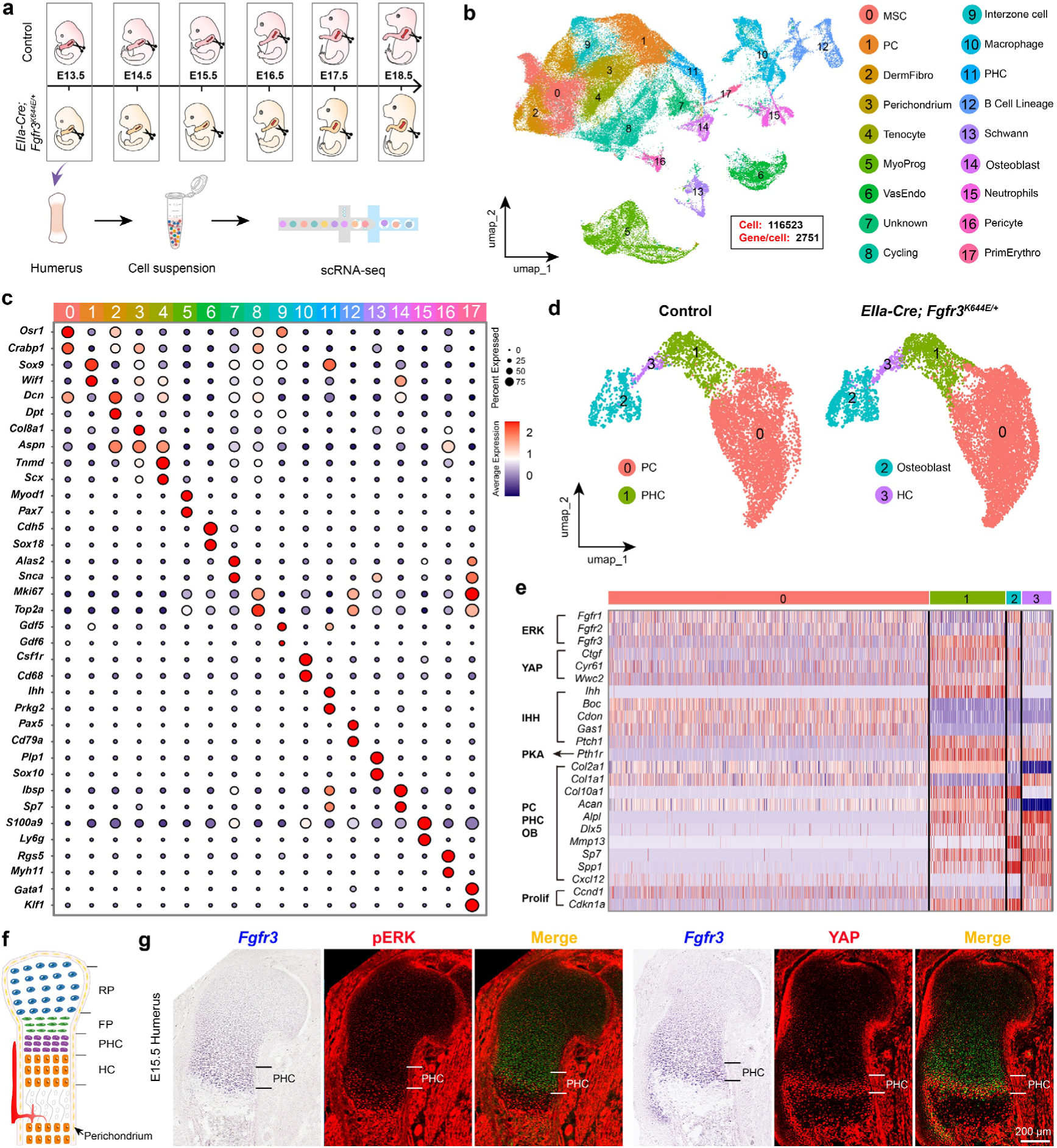
YAP signaling and FGFR3 exhibit the strongest activity in PHCs. **a, b,** scRNA-Seq of humeri from control and *Fgfr3-K644E* mice across 6 time points. **c,** Dotplots show marker gene expression across clusters. **d,** The chondrocyte lineage was isolated for further analysis. **e,** Heatmap of differentially expressed genes among PCs, PHCs, HCs, and osteoblasts (OBs) in control E16.5 humeri. **f,** Schematic of chondrogenesis. RP, round PCs; FP, flat PCs. **g,** Combined i*n situ* hybridization and immunostaining of E15.5 WT mouse proximal humeri reveal mutually exclusive expression patterns between *Fgfr3* and pERK, while *Fgfr3* and YAP exhibit largely overlapping expression.

Analysis of normal control humeri from E15.5, E16.5, and E17.5 revealed the following key patterns (Fig. 3e-g, and Extended Data Fig. 8): (1) *Fgfr1* and *Fgfr2* are expressed in PCs but severely downregulated in PHCs. (2) *Fgfr3* expression progressively increases in flat PCs, peaks in non-proliferating PHCs, and then declines in HCs—a pattern opposite to that of ERK signaling activity. (3) YAP signaling genes, *Cyr61* (*Ccn1*), *Ctgf* (*Ccn2*), and *Wwc2*, largely mirror the expression pattern of *Fgfr3*. (4) *Ihh* is expressed in PHCs and at lower levels in HCs, whereas its coreceptors (*Boc, Cdon, Gas1*) were almost exclusively expressed in PCs. (5) *Pth1r* expression was elevated in PHCs, HCs, and osteoblasts, thereby activating PTH1R-cAMP-PKA signaling ^3,5,23–25^, which in turn represses both IHH-SMO and YAP signaling ^18,26–29^ (Fig. 3e-g, and Extended Data Fig. 8). Finally, we also re-analyzed a published scRNA-Seq dataset of human femur and tibia at gestational week 9 ^30^, and found conserved expression patterns of these genes between human and mouse (Extended Data Fig. 9).

### *Fgfr3* mutation regulates ERK and YAP signaling to influence chondrocyte hypertrophy and ossification

Having analyzed chondrocytes in controls, we then examined corresponding populations in *Fgfr3-K644E* mice (Extended Data Fig. 10). In general, *Fgfr3-K644E* mutant PHCs displayed an immature state, as indicated by significant downregulation of genes typically enriched in PHCs, such as: *Acan, Alpl, Col10a1, Mef2c, Mef2d, Piezo1, Prkg1, Prkg2, Pth1r, Runx2, Sik3*, and *Tgfb2* (Extended Data Fig. 10a, b). In contrast, the osteoblast population exhibited a precociously differentiated state, characterized by upregulation of markers normally associated with terminal HCs and osteoblasts, including *Alpl, Bglap, Col1a1, Dmp1, Gja1, Mmp13, Pdgfd, Runx2, Slit2, Spp1*, and *Tnfrsf19* (Extended Data Fig. 10c, d). This polarized phenotype highlights the distinct temporal roles of FGFR3 in the developing humerus of *Fgfr3-K644E* mice.

We next separately analyzed gene expression changes between control and *Fgfr3*-*K644E* mutant mice at E15.5, E16.5, and E18.5 (Fig. 4). At E15.5, *Fgfr3-K644E* mutant PHCs delayed the onset of chondrocyte pre-hypertrophy, evidenced by the downregulation of characteristic PHC genes, including *Fgfr3* (*Fgfr3-K644E*) itself (Fig. 4a, d). This effect again stems from enhanced ERK signaling in mutant PHCs, as indicated by upregulation of ERK related genes such as *Etv1, Etv4, Etv5*, and *Spred1* in PHCs, and coincides with ERK-mediated repression of *Fgfr3* expression (Fig. 4d). Moreover, *Ccnd1* expression was upregulated, whereas that of the pro-differentiation factor *Cdkn1a* (*p21*) was downregulated. (Fig. 4d). Since PHCs are primarily post-mitotic, elevated ERK activity in PHCs of *Fgfr3-K644E* mice does not lead to bone overgrowth. Instead, it significantly delays the maturation of early PHCs.

**Fig. 4.**
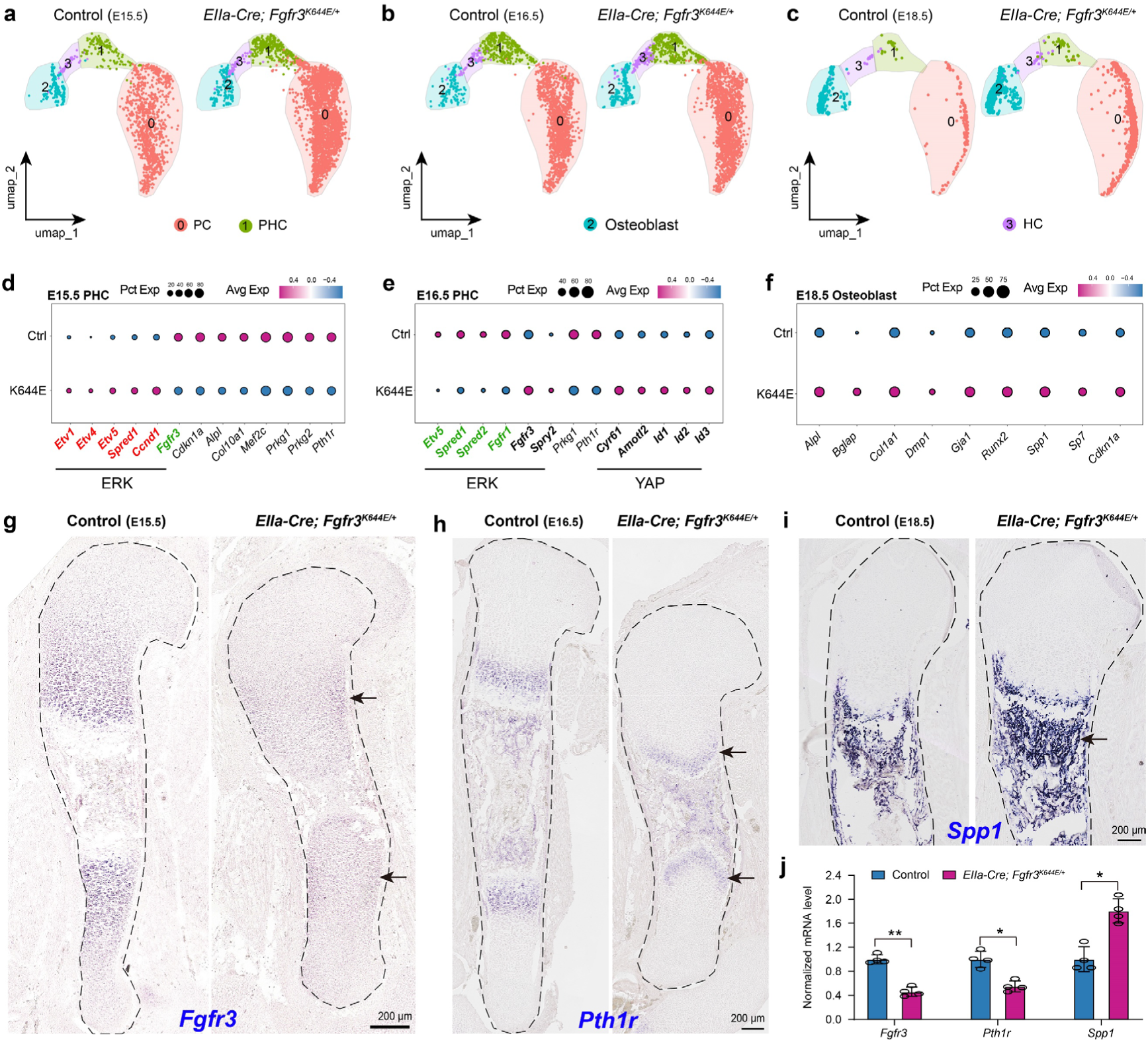
The *Fgfr3-K644E* mutation not only arrests PHC maturation but also induces premature humeral ossification. **a, d, g,** At E15.5, scRNA-Seq and *in situ* hybridization revealed that *Fgfr3-K644E* mutation upregulates ERK signaling and inhibits PHC pre-hypertrophy in humerus. *Fgfr3* expression was downregulated. **b, h, e,** At E16.5, *Fgfr3-K644E* mutation downregulated ERK and PTH1R-PKA signaling in PHCs, leading to the promotion of YAP target genes. The initiation of *Fgfr3* upregulation was noted. **c, f, I,** By E18.5, HCs and osteoblasts in *Fgfr3-K644E* mutants exhibited premature osteogenic differentiation. **j,** mRNA levels (*in situ* hybridization, arrows), were quantified at corresponding anatomical sites. n = 4; mean ± SEM; *, P<0.05; **, P<0.01 (unpaired Student’s *t-test*).

At E16.5, the *Fgfr3-K644E* mutation likely downregulated ERK signaling in PHCs, as shown by reduced expression of *Etv5*, *Spred1*, *Spred2*, and *Fgfr1* (Fig. 4b, e). Conversely, *Fgfr3* and *Spry2* expression was moderately elevated. Additionally, *Pth1r* and *Prkg1* remained downregulated in PHCs (Fig. 4e). The combined downregulation of ERK and PTH1R-PKA signaling strongly enhances YAP activity in PHCs, reflected by upregulation of YAP-associated genes such as *Cyr61, Amotl2, Id1, Id2*, and *Id3* (Fig. 4e). Enhanced YAP signaling promotes the nuclear translocation of unphosphorylated YAP/TAZ, which then bind within the nucleus to transcription factors such as TEAD, SMAD, and RUNX ^31–33^, thereby activating their target genes. Consistent with this mechanism, E18.5 *Fgfr3-K644E* mutant osteoblasts displayed a marked upregulation of mature osteoblast markers, including *Alpl, Bglap, Col1a1, Dmp1, Gja1, Runx2, Spp1, Sp7*, and *Cdkn1a* (Fig. 4c, f). Key findings were further validated by immunostaining and *in situ* hybridization (Fig. 4g-j; Extended Data Fig. 11–15). Thus, by E18.5, HCs and osteoblasts in *Fgfr3*-*K644E* mutants adopt a prematurely differentiated state that drives the accelerated maturation and ossification of bone.

## Discussion

This study reveals that the *Fgfr3-K644E* mutation acts through a unified self-limiting mechanism to cause both cortical overgrowth and skeletal restriction (Fig. 5a-d). Given that heightened FGF-FGFR-ERK signaling suppresses *Fgfr3* expression ^14,18,34–36^, *Fgfr3* expression can serve as an indicator of low ERK signaling activity in pERK expressing cells. Furthermore, as FGFR3 frequently exhibits greater sensitivity and responsiveness to FGF8/17/18 compared to FGFR1 and FGFR2 in many biological contexts ^9^, FGFR3 functions in *Fgfr3*-expressing cells to maintain ERK signaling at an appropriate level. Therefore, the oscillatory feedback between FGF-FGFR-ERK and FGFR3, alongside crosstalk with other signals, enables finely tuned regulation. Alterations in this dynamic balance during development, evolution, or disease significantly impact organ size and spatial cell fate in structures, including the cerebral cortex and skeleton. Indeed, we have recently identified a tripartite signaling network, marked by cross-repressive interactions among ERK/PKA, YAP/TAZ, and Hedgehog (HH) pathways, that orchestrates mammalian cortical development and evolution ^18^. In line with this principle, this study demonstrates that spatiotemporally regulated FGFR3 activity plays a prominent role in establishing the posterior-to-anterior developmental gradient of ependymal cells (Fig. 5b, c), a gradient phenomenon that has also been documented in earlier studies ^18,37,38^.

**Fig. 5.**
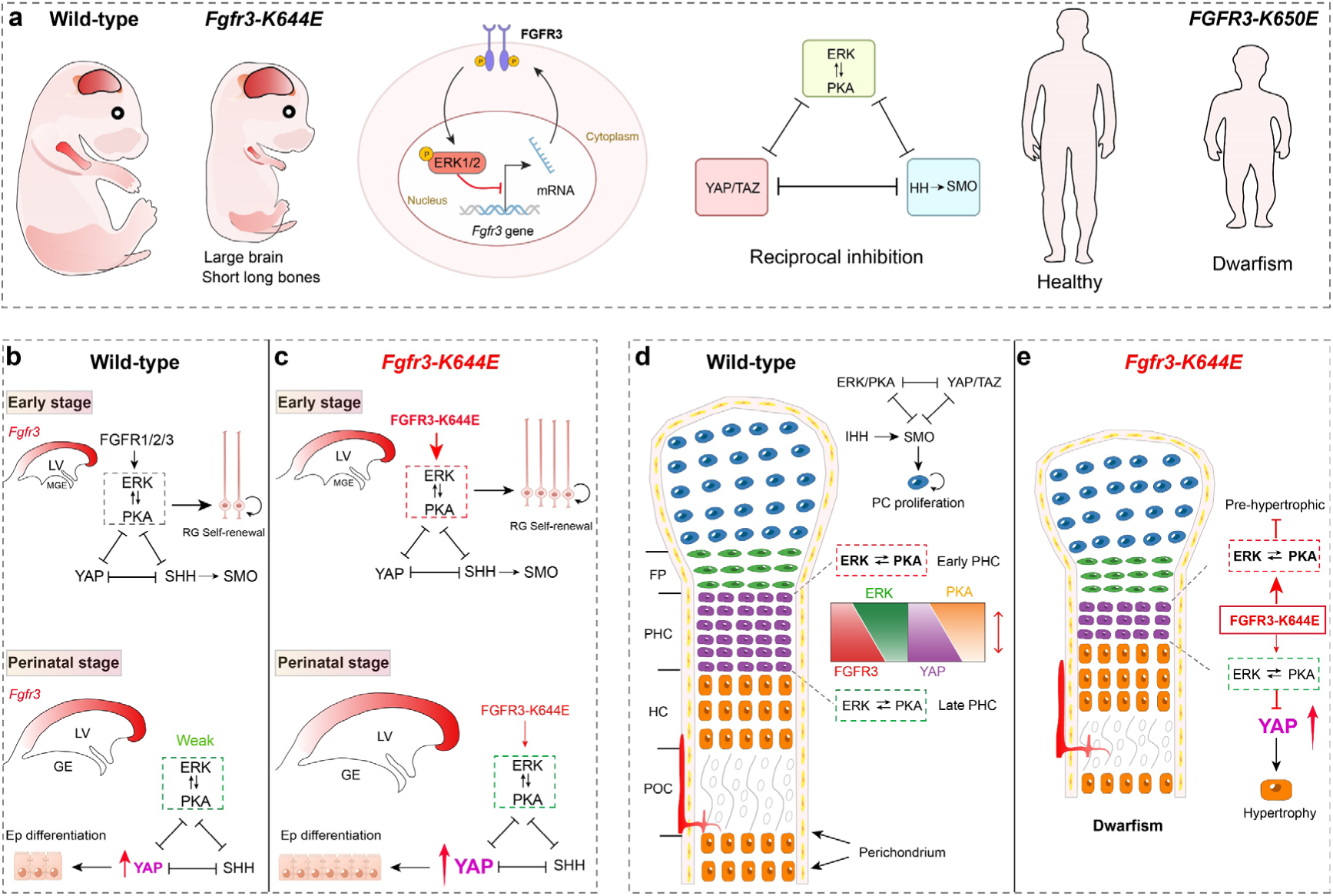
The pathogenic mechanism of *FGFR3-K644E* underlying macrocephaly-dwarfism. **a,** *Fgfr3-K644E* mutation engages a negative feedback loop, driving *Fgfr3* transcriptional repression which in turn downregulates ERK signaling. This mechanism establishes an oscillatory, self-limiting feedback circuit. The coordinated activity of integrated ERK/PKA, YAP, and HH signaling converges to drive cortical and long bone development, whose dysregulation underpins related disorders. **b,** High *Fgfr3* expression in caudal cortical RGs is a result of stronger ERK activity in the rostral cortex. This differential distribution ultimately establishes a posterior-to-anterior ependymal maturation gradient. **c,** The *Fgfr3-K644E* mutation expands caudal cortical RGs and prematurely polarizes ependymal maturation by initially activating ERK, then downregulating *Fgfr3*/ERK/PKA while enhancing YAP. **d,** Normal chondrogenesis relies on the coordinated interplay of the IHH-SMO, FGFR-ERK/PTH1R-PKA, and YAP/TAZ signaling pathways. IHH-SMO signaling promotes PC proliferation. *Fgfr3* expression is lowest in round PCs, moderate in flat PCs, highest in PHCs, and low in HCs—a pattern inversely correlated with ERK activity. In PHCs, the gradual reduction of ERK and PTH1R-PKA signaling results in YAP de-repression, nuclear accumulation, and eventual cellular hypertrophy. **e,** *Fgfr3* GOF mutations disrupt this balance through a biphasic mechanism: by initially inhibiting pre-hypertrophy/hypertrophy via ERK hyperactivation, and subsequently unleashing YAP-driven hypertrophy through marked downregulation of ERK/PKA signaling, ultimately resulting in premature ossification and dwarfism.

We propose that the tripartite network similarly governs bone development and growth, following conserved mechanistic principles (Fig. 5a, d, e). In PCs, IHH-SMO signaling is the primary driver of proliferation during normal long bone development ^39^, and concurrently represses ERK/PKA and YAP/TAZ signaling ^18,26–29^. In early PHCs adjacent to PCs, mildly active ERK signaling synergizes with PTH1R-PKA to suppress IHH-SMO and prevent premature YAP activation. In contrast, in distal PHCs farther from PCs, downregulated ERK signaling, combined with reduced PTHLH-mediated PTH1R-PKA activity, results in decreased PKA activity, which in turn promotes YAP activation (Fig. 5d). The Hippo-YAP pathway regulates multiple aspects of chondrogenesis—including cell polarity, density, morphology, and mechanosensation—and drives substantial chondrocyte volume expansion, which increases roughly 20-fold during this process ^5,32,40–44^. In *Fgfr3* GOF mutations, early ERK hyperactivation impedes normal PHC maturation, while later ERK downregulation further accelerates PKA downregulation and enhances YAP signaling. Collectively, these disruptions lead to accelerated depletion of PHCs, reduced PC numbers, premature hypertrophy and mineralization of HCs/osteoblasts, and ultimately result in dwarfism (Fig. 5e). Excessive YAP signaling is known to cause chondrodysplasia and dwarfism ^41,42^. PTH1R-PKA signaling dysfunction leads to chondrodysplasia or accelerated chondrocyte differentiation ^23–25,45^, potentially also through misregulated YAP signaling. Therefore, the ERK, PKA, HH-SMO, and YAP/TAZ signaling pathways collectively establish an integrative model of mutual inhibition, which underlies both normal development and disease. This network reveals novel therapeutic targets for skeletal and neurological disorders.

## Acknowledgments

We thank Y. You, and Z. Li for technical assistance.

## Author contributions

Conceptualization: Z.Y. Data curation: Z.Z. and Z.X. Funding acquisition: Z.Z., Z.X., and Z.Y. Investigation: Z.Z., Z.X., T.F., W.Z., Z.S., C.Y., F.Y., J.L., and J.D. Resources: Z.Y. Supervision: Z.Y. Writing: Z.Y. All of the authors contributed to reviewing and editing of the manuscript.

## Data Availability Statement

The datasets (scRNA-Seq from 20 samples, Supplementary Table 3) generated during the current study are available in the Gene Expression Omnibus (GEO: GSE316124). scRNA-Seq data from human femur and tibia at gestational week 9, are available via the European Nucleotide Archive via accession number PRJEB28278 9 ^30^.

## Additional information

Supporting Information is available from the Online.

## Methods

### Mice

All procedures involving animals were approved by and performed in accordance with the guidelines of the Fudan University Shanghai Medical College Animal Ethics Committee (No. 20230301-141). *EIIa-Cre* (JAX no. 003724) ^46^, *Emx1-Cre* (JAX no. 005628) ^47^, *Nes-Cre* (JAX no. 003771) ^48^, and *Rosa^MEK1DD^* (JAX no. 012352) ^49^, were described previously. *Rosa^MEK1DD^* mice, which harbor a Cre-inducible, constitutively active rat *Map2k1* allele (S218D/S222D) and an EGFP reporter, were crossed with *Emx1-Cre* or *Nes-Cre* drivers to establish brain tissue-specific systems enabling controlled, sustained ERK signaling activation. The *Fgfr3-K644E* knock-in mutant mice used in this study were generated by GemPharmatech Co., Ltd. Nanjing, China. The targeted knock-in strategy was employed to generate a conditional mouse model for the *Fgfr3-K644E* mutation. The design involved inserting a mutant exon 14 (carrying the K644E point mutation) into the genomic locus between introns 9 and 10 of the mouse *Fgfr3* gene. To enable conditional control, a floxed cassette containing exons 10–18 (covering the critical tyrosine kinase domain and C-terminal coding sequence) along with a transcriptional stop signal was introduced upstream of the mutant exon. At the protein level, this mutation results in the substitution of lysine (K) with glutamic acid (E) at residue 644, driven by a single-nucleotide change from AAG to GAG. This amino acid alteration constitutively activates the FGFR3 receptor and mimics human skeletal dysplasia–associated mutations. The conditional design allows tissue- or time-specific expression of the mutant allele upon Cre-mediated recombination, thereby offering a precise tool to study the spatiotemporal roles of hyperactive FGFR3 signaling in development and disease. *Fgfr3-K644E* mice are available upon request. Control mice consisted of both wild-type (WT) littermates and Cre-negative littermates from the same crosses, ensuring that all groups shared an equivalent genetic background aside from the experimental manipulation. The day of detecting a vaginal plug was designated as E0.5. The day of birth was designated as P0. The sexes of the embryonic and early postnatal mice were not determined. For immunostaining analysis, normally, brains and humeri from 2-5 independent experiments were processed (n values refer to numbers of samples analyzed).

### Tissue preparation

Embryos were isolated from deeply anesthetized pregnant mice. Brains were dissected and fixed overnight in 4% paraformaldehyde (PFA) pre-treated with diethylpyrocarbonate (DEPC). Postnatal mice were deeply anesthetized and transcardially perfused with phosphate-buffered saline (PBS), followed by 4% PFA. All brains were post-fixed overnight in 4% PFA at 4°C and subsequently dehydrated in 30% sucrose for at least 24 hours. The brains were then embedded in O.C.T. compound (Sakura Finetek) and stored at -80°C. For analysis, mouse brains were sectioned into 20-μm-thick slices.

Skeletal preparations were performed using a modified standard protocol for cartilage and bone staining ^50–52^. Briefly, specimens were fixed in 99% ethanol for 24 h, followed by dehydration in acetone for an additional 24 h. The samples were then sequentially stained with Alcian Blue (Sigma A-3157) for cartilage and Alizarin Red (0.05 mg/mL in 0.5 % KOH) for bone, each step performed at room temperature with gentle agitation. After staining, samples were thoroughly rinsed in distilled water and transferred into a clearing solution consisting of 1 % potassium hydroxide supplemented with 20 % glycerol, and incubated at 37 °C overnight. Clearing was continued at room temperature until the soft tissues became transparent and the skeletal structures were clearly visible. For long-term preservation, specimens were progressively transferred into graded glycerol solutions (50 %, 80 %, and finally 100 % glycerol) for stable storage and imaging. For histological analysis, samples were fixed in 4% PFA in phosphate-buffered saline (PBS) at 4°C overnight, followed by cryoprotection through immersion in 30% sucrose at 4°C for at least 24 h or until the tissue sank. The samples were then embedded in optimal cutting temperature (O.C.T.) compound (Sakura Finetek), frozen on dry ice, and stored at –80°C until sectioning. Serial sections of long bones were cut at thicknesses of 12 μm or 20 μm using a cryostat and mounted on glass slides for subsequent staining and analysis.

### FlashTag labeling

FlashTag labeling was performed via in utero injection or in neonatal mice, followed by fluorescence-activated cell sorting (FACS) to isolate live labeled cells ^53,54^. For embryonic labeling, a volume of 0.5 μL of FlashTag–CellTrace Yellow solution (Life Technologies, #C34567; 10 mM) was injected into the lateral ventricles of control and *Fgfr3-K644E* mouse embryos at embryonic day 17.5 (E17.5). Labeled cortical progenitors were collected and analyzed 24 h later at E18.5. For postnatal labeling, the same CellTrace Yellow solution was injected into the lateral ventricles of neonatal controls and mutant mice at postnatal day 0 (P0), with analysis performed P2 (*Emx1-Cre; Fgfr3^K644E/K644E^*). FlashTag labeled mice were euthanized and brains were rapidly dissected and placed in ice-cold Hanks’ Balanced Salt Solution (HBSS; Gibco, #14175-095). Cortical tissues were microdissected, enzymatically and mechanically dissociated into single-cell suspensions, and subjected to FACS to purify FlashTag-positive cells. Finally, Single-Cell RNA Sequencing (scRNA-Seq) libraries were prepared from the FACS-sorted cells using the 10X Genomics Chromium platform and sequenced according to the manufacturer’s protocol.

### Immunohistochemistry

Immunofluorescence staining was conducted on 10 μm or 20 μm cryosections of mouse brains or humeri following standard procedures. Sections were first rinsed in Tris-buffered saline (TBS; 0.01 M Tris–HCl, 0.9% NaCl, pH 7.4) for 10 min, then permeabilized with 0.5% Triton X-100 in TBS for 30 min at room temperature (RT). Subsequently, sections were blocked in a solution containing 5% donkey serum and 0.5% Triton X-100 in TBS (pH 7.2) for 2 h at RT. After blocking, sections were incubated overnight at 4 °C with primary antibodies diluted in blocking buffer. The following day, sections were washed three times with TBS (10 min per wash) and incubated with species-matched fluorescent secondary antibodies (Jackson ImmunoResearch; 1:500 dilution) for 2 h at RT in the dark. Following three additional TBS washes (10 min each), nuclei were counterstained with 4′,6-diamidino-2-phenylindole (DAPI; Sigma, 200 ng/mL in TBS) for 2 min. All primary antibodies used in this study, including supplier, catalog number, host species, and dilution, are listed in Supplementary Table 1.

### Western Blotting

Cortices from E12.0 or E12.5 mouse embryos were dissected under a stereomicroscope and lysed in RIPA Lysis Buffer (Beyotime, #P0013K; containing 50 mM Tris pH 7.4, 150 mM NaCl, 1% Triton X-100, 1% sodium deoxycholate, 0.1% SDS, and 1% PMSF [Beyond, #ST506-2]). Protein concentration was measured using a BCA assay kit (EpiZyme, #ZJ101). Equal amounts of protein were separated on 8% SDS-PAGE gels and transferred to PVDF membranes. The membranes were blocked in 5% non-fat milk for 1 h at room temperature and then incubated overnight at 4 °C with the following primary antibodies diluted in blocking buffer: mouse anti-β-Tubulin (1:3000, Abmart, #M20005F); rabbit anti-HOPX (1:2000, Oasis Biofarm, #OB-PRT015); rabbit anti-pERK1/2 (1:4000, Cell Signaling Technology, #4370). After three washes with TBST, membranes were incubated for 1 h at room temperature with horseradish peroxidase (HRP)-conjugated secondary antibodies: Goat Anti-Mouse IgG (H+L) (1:3000, Beyotime, #A0216) and Goat Anti-Rabbit IgG (H+L) (1:3000, Beyotime, #A0208). Following three additional TBST washes, protein bands were visualized using BeyoECL Star detection reagent and imaged with an e-blot biomolecular imager. Band intensities were quantified using ImageJ software.

### scRNA-Seq

Mouse cortical or humeral tissues were dissected and enzymatically/mechanically dissociated into single-cell suspensions in cold PBS containing 0.04% bovine serum albumin. Cell viability and concentration were assessed using trypan blue exclusion. For selected samples (E18.5 and P1 cortices), FlashTag labeled live cells were further enriched by fluorescence-activated cell sorting (FACS) prior to library preparation. Single-cell libraries were constructed using the Chromium Next GEM Single Cell 3′ Kit v3.1 (10X Genomics) on the Chromium Controller according to the manufacturer’s instructions (document CG00052 Rev C). Purified cDNA libraries were quantified with an Agilent 2100 Bioanalyzer High Sensitivity DNA Assay and sequenced on an Illumina NovaSeq 6000 platform using a 150-bp paired-end configuration, aiming for a median depth of >50,000 reads per cell. In total, this study generated and systematically characterized 20 novel mouse scRNA-Seq datasets. Complete metadata, including tissue source, developmental stage, genotype, are summarized in Supplementary Table 2.

A published human fetal bone scRNA-Seq dataset was reanalyzed in this study ^30^. The dataset was derived from the developing proximal and distal regions of the femur and tibia of a human fetus at gestational week 9 (GW9; approximately post-conception week 7, PCW7). Original experimental details are described in the respective publication ^30^.

### scRNA-Seq data analysis

scRNA-Seq reads were aligned to the mm10 reference genome and quantified using ‘cellranger count’ (10x Genomics, v.7.1.0). Count data was further processed using the ‘Seurat’ R package (v.5.1.0). Different quality control (QC) thresholds were applied across batches to account for batch-specific differences in sequencing depth and data quality. Initial clustering was performed independently for each sample using Seurat to identify potential low-quality cell clusters. Low-quality clusters were defined as those enriched for cells with a low number of unique molecular identifiers (UMIs) and/or a high proportion of mitochondrial gene-derived UMIs. The resulting filtered count matrices were subsequently analyzed using the DoubletFinder (v2.0) package to estimate doublet scores for each cell in a sample-wise manner. Cell clusters showing a high enrichment of predicted doublets, as indicated by elevated doublet scores, were considered doublet clusters and removed from downstream analyses.

After QC filtering and doublet removal, gene expression data were normalized using Seurat’s global-scaling normalization method (“LogNormalize”) with default parameters. The top 3,000 highly variable genes were identified using the FindVariableFeatures function with the “vst” method. Principal component analysis (PCA) was then performed, and the top 50 principal components (PCs), selected based on variance explained, were retained for downstream neighborhood graph construction and clustering using the FindNeighbors and FindClusters functions. Uniform Manifold Approximation and Projection (UMAP) was applied for low-dimensional visualization of cell clusters. Differentially expressed genes (DEGs) among clusters were identified using the Wilcoxon rank-sum test as implemented in Seurat, and *P* values were adjusted for multiple testing using the Benjamini– Hochberg false discovery rate (FDR) correction.

### mRNA *in situ* hybridization

Mouse mRNA *in situ* hybridization was performed on 20 μm cryosections using digoxigenin (DIG)-labeled riboprobes. We conducted *in situ* hybridization with at least two replicates for each time point. Sections were post-fixed in 4% paraformaldehyde (PFA) for 20 min prior to hybridization. The ISH procedure followed a previously described protocol ^55^. Riboprobes were synthesized from DNA templates cloned from mouse fetal cortical and long bone cDNA. Total RNA was extracted from mouse cortex and long bone tissues using the RNeasy Kit (Qiagen), reverse-transcribed with Superscript III Reverse Transcriptase (Invitrogen) and random hexamers, and used as the template for PCR amplification. Gene-specific primers were designed with Primer3 and PCR was carried out using Phusion High-Fidelity DNA Polymerase (Thermo Scientific). Riboprobes were made from mouse cDNAs amplified by PCR using the primers in Supplementary Table 3.

### Image acquisition and analysis

All brain section images in this study were acquired using an Olympus VS200 Automated Slide Scanner (10X, 20X) or an FV3000 confocal microscope system (40X). Images were processed using Adobe Photoshop for clarity, false colorization, and overlay as needed. Both Adobe Photoshop and Adobe Illustrator were used to adjust images without altering the original data.

To analyze the spatial relationship between *Fgfr3* and ERK, as well as between *Fgfr3* and YAP, we performed digital image superimposition. Specifically, bright-field in situ hybridization images for mRNA localization (*Fgfr3*) and immunofluorescence images for protein detection (ERK and YAP/TAZ) were acquired from serial adjacent humerus sections. These paired images were then precisely aligned and digitally fused using Adobe Photoshop to enable direct visual comparison and assessment of co-localization patterns. Combined in situ hybridization and immunostaining analysis of E15.5 WT mouse proximal humeri shows that *Fgfr3* and pERK exhibit mutually exclusive expression patterns, with *Fgfr3* and YAP/TAZ, in contrast, displaying substantial co-localization (see Fig. 3g).

### Quantification and statistical analysis

Quantitative images were acquired using the Olympus VS200 Automated Slide Scanner (10X or 20X). Statistical analyses were performed using GraphPad Prism 10 and IBM SPSS Statistics 29.0. For each experiment, at least four control or mutant samples were analyzed. The following quantifications were performed:

1. FOXJ1-positive cells were quantified in the P2 cortical VZ/SVZ of Control, *Emx1-Cre; Fgfr3^K644E/+^*, and *Emx1-Cre; Fgfr3^K644E/K644E^* mice. Four corresponding coronal sections at the same rostro-caudal level were selected from each brain for analysis. (Fig. 2h).
2. The mRNA signal intensities of *Fgfr3* (E15.5), *Pth1r* (E16.5), and *Spp1* (E18.5) were quantified in four humerus sections from 2 mice (in situ hybridization was performed in duplicate with 2 mice serving as biological replicates) using ImageJ software. Values were normalized to the mean intensity of the Control group, and statistical analyses were performed using GraphPad Prism 10 (Fig. 4j).
3. The area of the P0 hemisphere (delineated by dashed lines in Extended Data Fig.1C) was quantified using Adobe Photoshop (n = 4 mice per group), and data were analyzed using GraphPad Prism 10 (Extended Data Fig.1D). Similarly, the humerus length at P0 (n = 4 mice per group) was measured using Adobe Photoshop. Lengths were normalized to the mean of the Control group and analyzed using GraphPad Prism 10 (Extended Data Fig. 1f).
4. HOPX immunofluorescence intensity was quantified in the cortical VZ/SVZ of E18.5 Control and *Emx1-Cre; Fgfr3^K644E/+^* mice. For each animal, 4–6 coronal sections were analyzed using ImageJ software (*n* = 4 mice per group). Signal intensities were normalized to the mean of the Control group, and statistical analyses were performed using GraphPad Prism 10 (Extended Data Fig. 5f).
5. The cortical area of the hemisphere (delineated by dashed lines in Extended Data Fig.6A, D) was measured in P27 Control, *Emx1-Cre; Fgfr3^K644E/+^*, and *Emx1-Cre; Rosa^MEK1DD/+^*mice using Adobe Photoshop (*n* = 4 mice per group). Values were normalized to the mean of the Control group (set to 1) to calculate relative cortical areas. Statistical analyses were performed using GraphPad Prism 10 (Extended Data Fig. 6b).
6. The cortical areas of the hemispheres (delineated by dashed lines in Extended Data Fig.7A and E) were quantified using Adobe Photoshop for P47 *Nestin-Cre; Fgfr3^K644E/+^* and P6 *Nestin-Cre; Rosa^MEK1DD/+^* cohorts, along with their respective Controls (*n* = 4 mice per group). Values were normalized to the mean of the corresponding Control group (set to 1) and analyzed using GraphPad Prism 10 (Extended Data Fig. 7b, f).

Data are presented as mean ± SEM (standard error of the mean) and were plotted using GraphPad Prism. For statistical comparisons, we used either one-way ANOVA (with Tukey’s post-hoc test) or an unpaired Student’s t-test, depending on experimental design.

---

**Extended Data Fig. 1.**
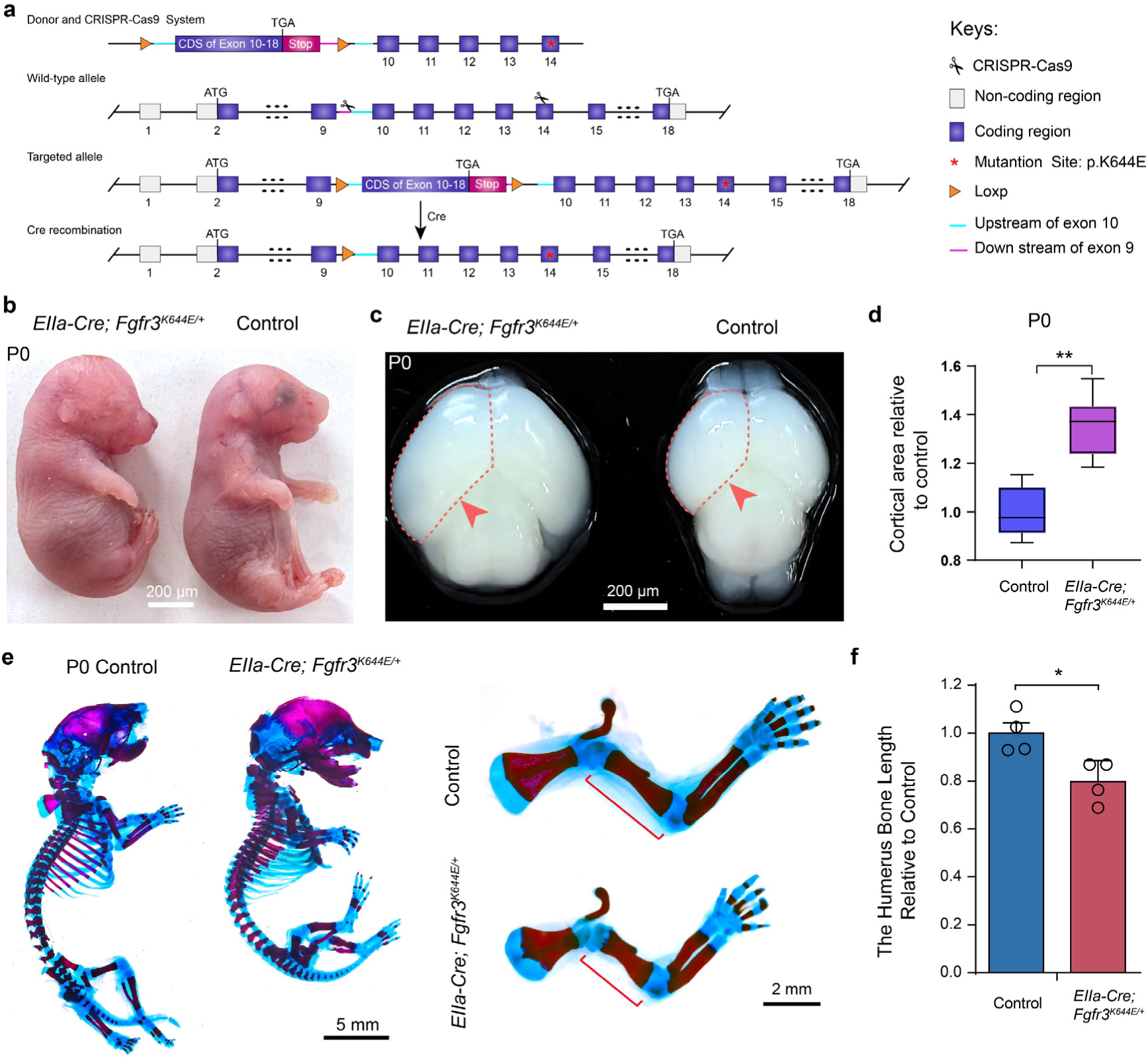
The *Fgfr3-K644E* (*EIIa-Cre; Fgfr3^K644E/+^*) mouse is a gain-of-function (GOF) model that phenocopies human Thanatophoric Dysplasia type II (TDII). **a.** Strategy for generating a conditional *Fgfr3^K644E/+^* knock-in mouse model. A mutant exon 14 (carrying the *Fgfr3-K644E* mutation) was inserted between introns 9 and 10 of the mouse *Fgfr3* gene, together with a loxP-exon 10–exon 18 CDS–stop–loxP cassette. This mutation substitutes lysine (K) at position 644 with glutamic acid (E), corresponding to a codon change from AAG to GAG. **b-f.** The *Fgfr3-K644E* mouse model phenocopies human TDII. The mutant mice are neonatal lethal (**b**), exhibit an enlarged cerebral cortex (**c, d**), and display shortened long bones and dwarfism (**e, f**). **e.** Alcian Blue (cartilage) / Alizarin Red (bone) staining at P0. (**d, f**) Quantification of the cortical area and humerus bone length ratios in *Fgfr3-K644E* mice relative to controls. n = 4; mean ± SEM; *P <0.05, **P <0.01; unpaired Student’s *t-test*.

**Extended Data Fig. 2.**
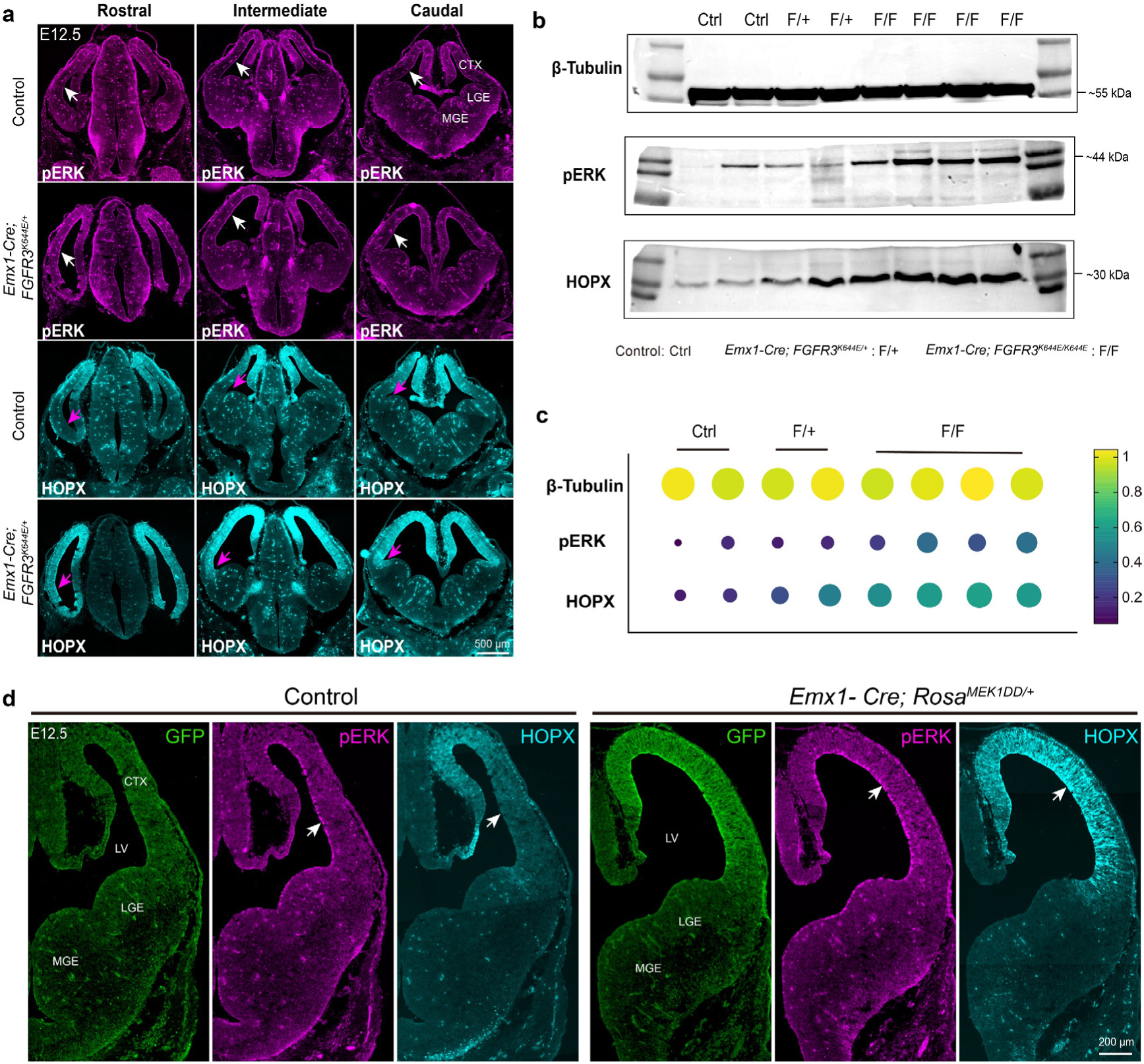
*Emx1-Cre; Fgfr3^K644E/+^* mice exhibited cortical phenotypes at E12.5 that were similar to those observed in *Fgfr3-K644E* mutant mice. **a.** Immunostaining revealed a subtle increase in pERK and strong *HOPX* expression in E12.5 cortical RGs of *Emx1-Cre; Fgfr3^K644E/+^* mice compared to controls (arrows). (**b, c**) Western blotting revealed a modest increase in pERK and strong *HOPX* expression in E12.5 cortices of *Emx1-Cre; Fgfr3^K644E/+^* and *Emx1-Cre; Fgfr3^K644E/K644E^* mice compared to controls. **d.** E12.5 cortical RGs in *Emx1-Cre; Rosa^MEK1DD/+^* mice showed a significant increase in both pERK and HOPX protein levels (indicated by arrows). CTX, cortex; LV, lateral ventricle; LGE, lateral ganglionic eminence; MGE, medial ganglionic eminence

**Extended Data Fig. 3.**
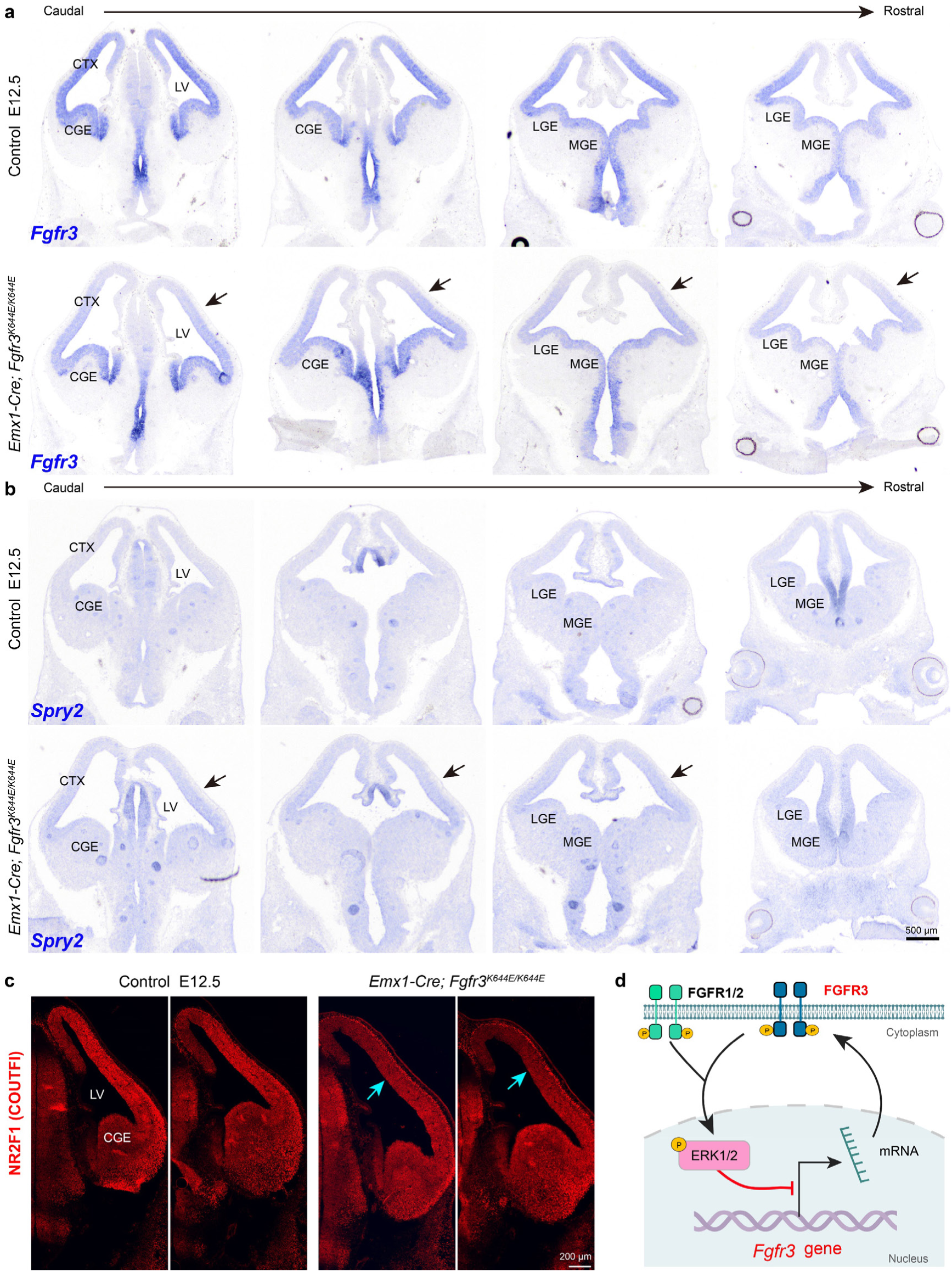
FGFR3-K644E suppresses endogenous *Fgfr3* expression in cortical RGs at E12.5. **a-c.** In situ hybridization and immunostaining revealed a modest upregulation of *Spry2* alongside downregulation of *Fgfr3* and NR2F1 (COUP-TFI) in E12.5 cortical RGs of *Emx1-Cre; Fgfr3^K644E/K644E^* mice compared to controls (indicated by arrows). **d**. *Fgfr3-K644E* mutation induces a negative feedback loop, downregulating its own transcription via ERK signaling. Similarly, in cortical RGs, FGF-FGFR1/2/3-ERK signaling triggers a feedback mechanism that downregulates ERK through *Fgfr3*.

**Extended Data Fig. 4.**
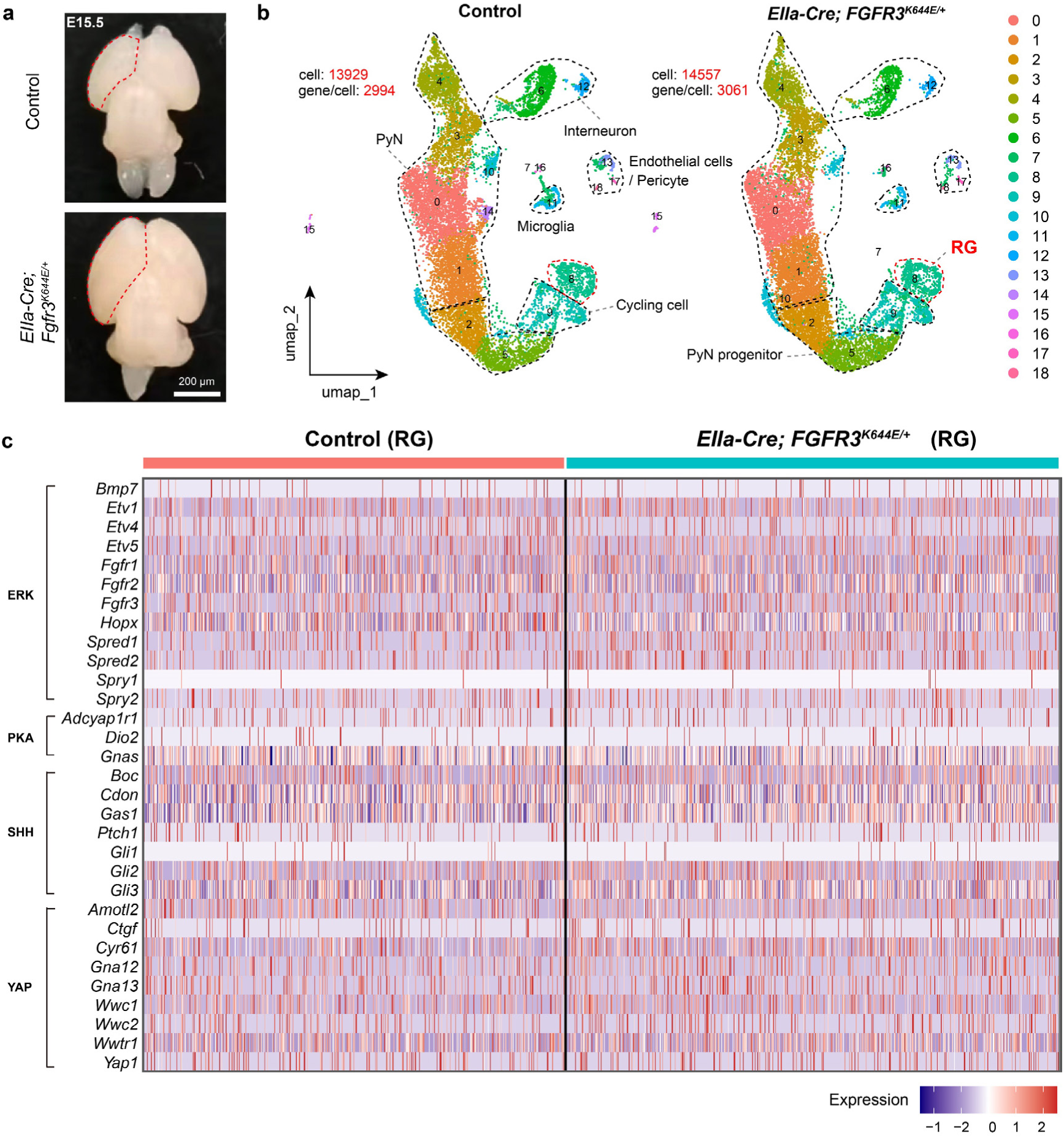
The *Fgfr3-K644E* mutant cortical RGs show no significant change in ERK signaling at E15.5. **a-c.** Despite the visibly enlarged cortex in *Fgfr3-K644E* mutants at E15.5, scRNA-Seq analysis detected no significant alterations in ERK, PKA, SHH, or YAP signaling within cortical RGs (neural stem cells, cluster 8). Note that *Fgfr1* and *Fgfr2* are expressed in cortical RGs and may buffer fluctuations in ERK signaling.

**Extended Data Fig. 5.**
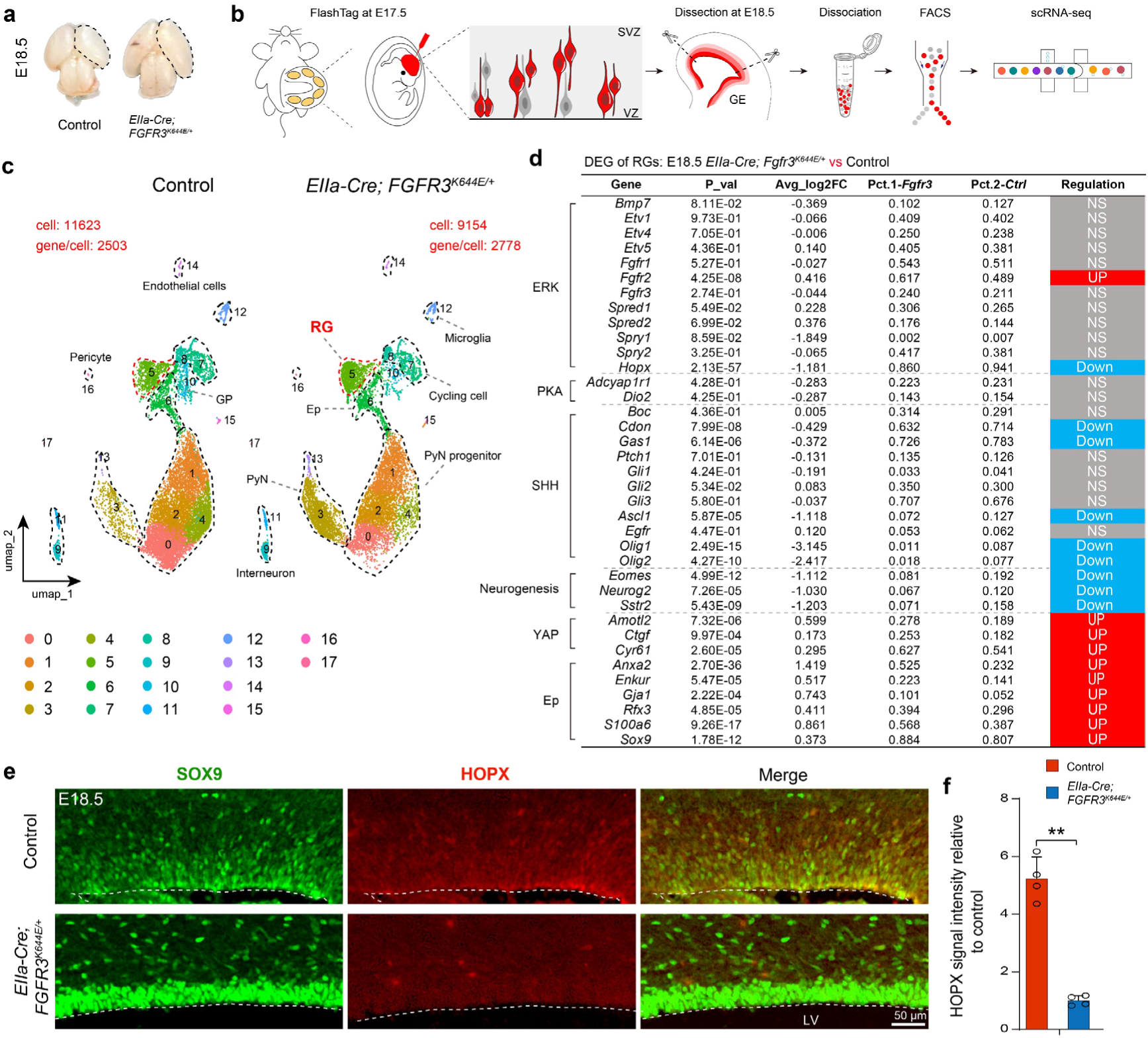
The *Fgfr3-K644E* mutation downregulates ERK, PKA, and SHH signaling but concurrently activates YAP signaling (YAP/TAZ activity) in cortical RGs at E18.5. **a, b.** Cortical progenitor cells were labeled and enriched from littermate control (WT) and *Fgfr3-K644E* mutant brains for scRNA-Seq analysis. FlashTag-CellTrace Yellow was injected into the lateral ventricle at E17.5, and FlashTag-positive cortical cells were isolated by fluorescence-activated cell sorting (FACS) at E18.5. **c.** Annotation of cortical cell clusters. **d.** scRNA-Seq analysis revealed differentially expressed genes (DEGs) between the cortical RGs (cluster 5) of *Fgfr3-K644E* mutants and their WT littermate controls at E18.5. Notably, cortical RGs exhibited a significant downregulation of *Hopx* expression. **e.** Immunostaining showed that HOPX expression was markedly downregulated in cortical RGs at E18.5, while SOX9 expression was increased. **f.** Quantification of HOPX expression levels in mutant cortical RGs compared with controls. n = 4; mean ± SEM; **P <0.01, (unpaired Student’s *t-test*).

**Extended Data Fig. 6.**
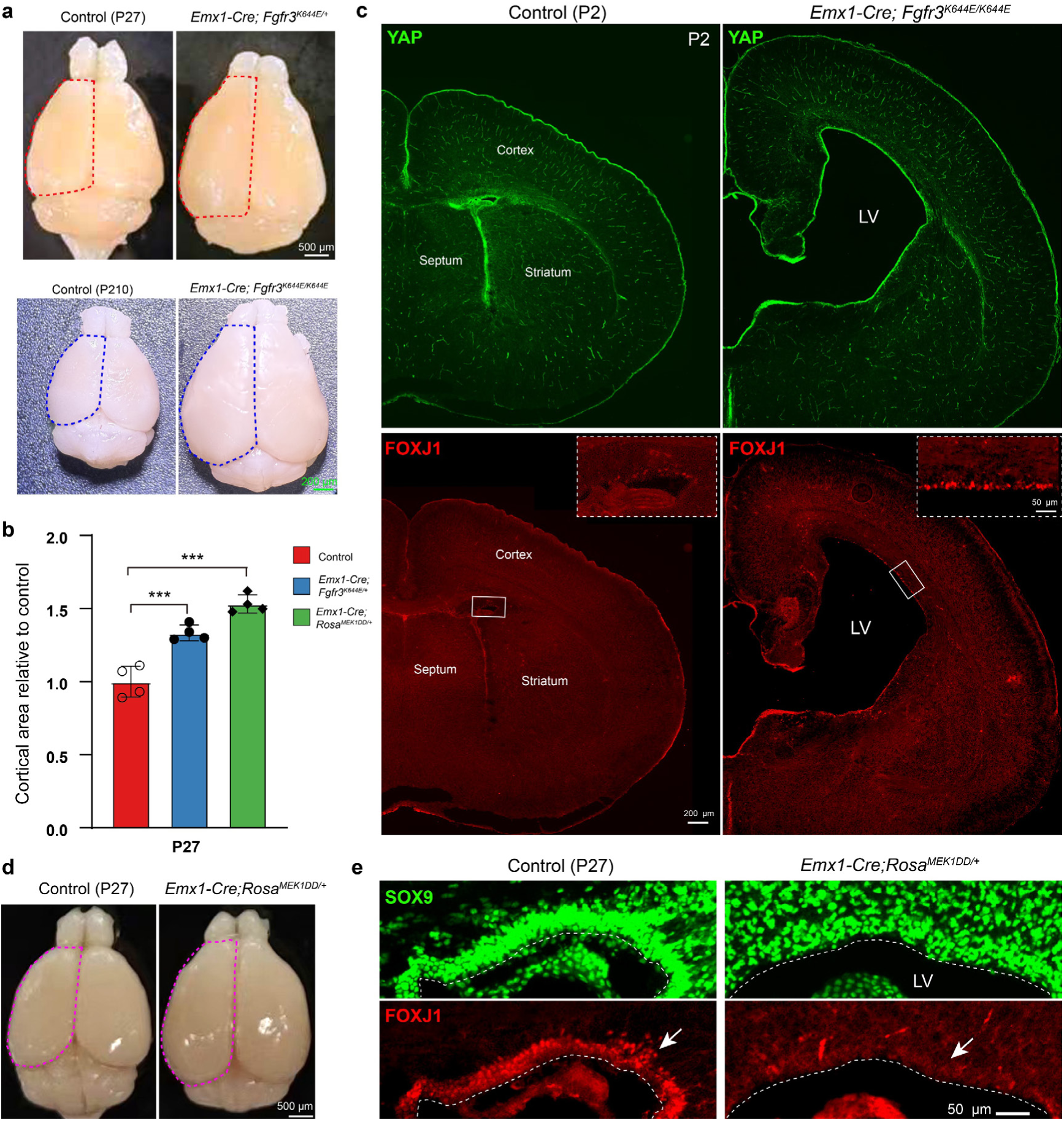
Although both *Emx1-Cre; Fgfr3^K644E/+^* and *Emx1-Cre; Rosa^MEK1DD/+^*mice exhibit cortical enlargement, they lead to distinct phenotypic outcomes in ependymal cells. **a.** In *Emx1-IRES-Cre* knock-in mice, Cre recombinase expression is driven by the endogenous *Emx1* locus starting from E10.5 in cortical neuroepithelial cells. *Emx1-Cre; Fgfr3^K644E/+^* mice survived long-term. **b.** Quantification revealed a larger cortical area in *Emx1-Cre; Fgfr3^K644E/+^* and *Emx1-Cre; Rosa^MEK1DD^* mice at P27 (n = 4; mean ± SEM; ***P <0.001; one-way ANOVA and Tukey’s post-hoc test). **c.** *Emx1-Cre; Fgfr3^K644E/K644E^* mice exhibited premature differentiation of cortical ependymal cells (FOXJ1⁺ cells) compared with controls at P2. Note that YAP was strongly expressed in the mutant cortical ventricular zone (VZ), with YAP/TAZ localized to the nucleus. **d, e.** In contrast, *Emx1-Cre; Rosa^MEK1DD/+^* mice showed defective development of cortical ependymal cells, characterized by a loss of FOXJ1⁺ cells. *Emx1-Cre; Rosa^MEK1DD/+^* mice died within three months of age.

**Extended Data Fig. 7.**
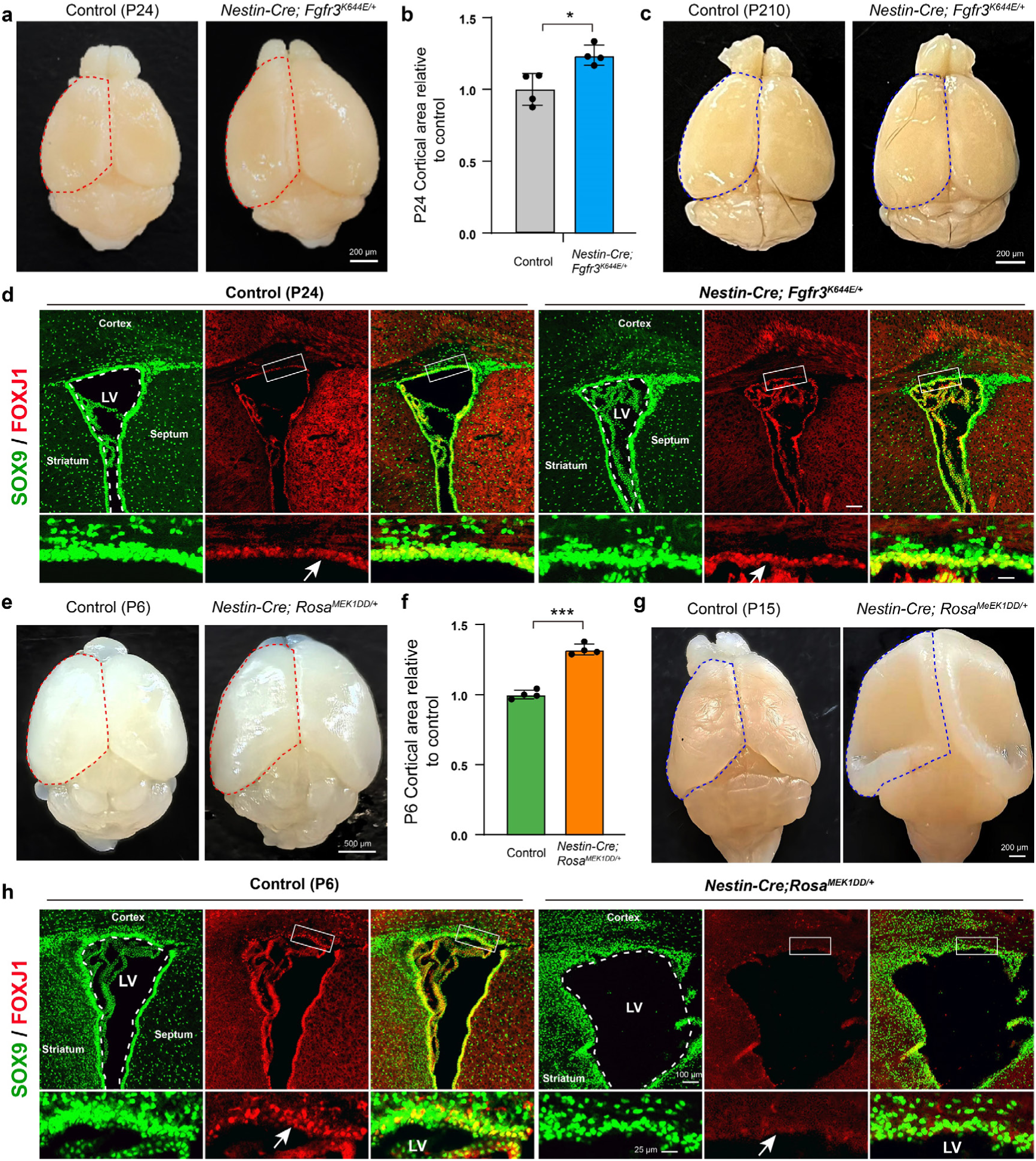
Both *Nes-Cre; Fgfr3^K644E/+^* and *Nes-Cre; Rosa^MEK1DD/+^* mice exhibit cortical enlargement, while their effects on ependymal cell phenotypes diverge. **a-d.** *Nes-Cre* (*Nestin-Cre*) is expressed in forebrain RGs from ∼E11.5. *Nes-Cre; Fgfr3^K644E/+^* mice survived long term and displayed essentially normal development of ependymal cells (FOXJ1⁺ cells). Quantification revealed a larger cortical area in *Nes-Cre; Fgfr3^K644E/+^* mice at P24 (n = 4; mean ± SEM; *P <0.05, unpaired Student’s *t-test*) (**b**). (**e-h**) In contrast, *Nes-Cre; Rosa^MEK1DD/+^* mice exhibited defective ependymal development, marked by loss of FOXJ1⁺ cells, severe hydrocephalus, and death within three weeks of age. Quantification revealed a larger cortical area in *Nes-Cre; Rosa^MEK1DD/+^*mice at P6 (n = 4; mean ± SEM; ***P <0.001, unpaired Student’s *t-test*) (**f**). Note that a subset of *Nes-Cre; Rosa^MEK1DD/+^* mice presented with hydrocephalus and a markedly thinned cerebral cortex as early as P15 (**g**).

**Extended Data Fig. 8.**
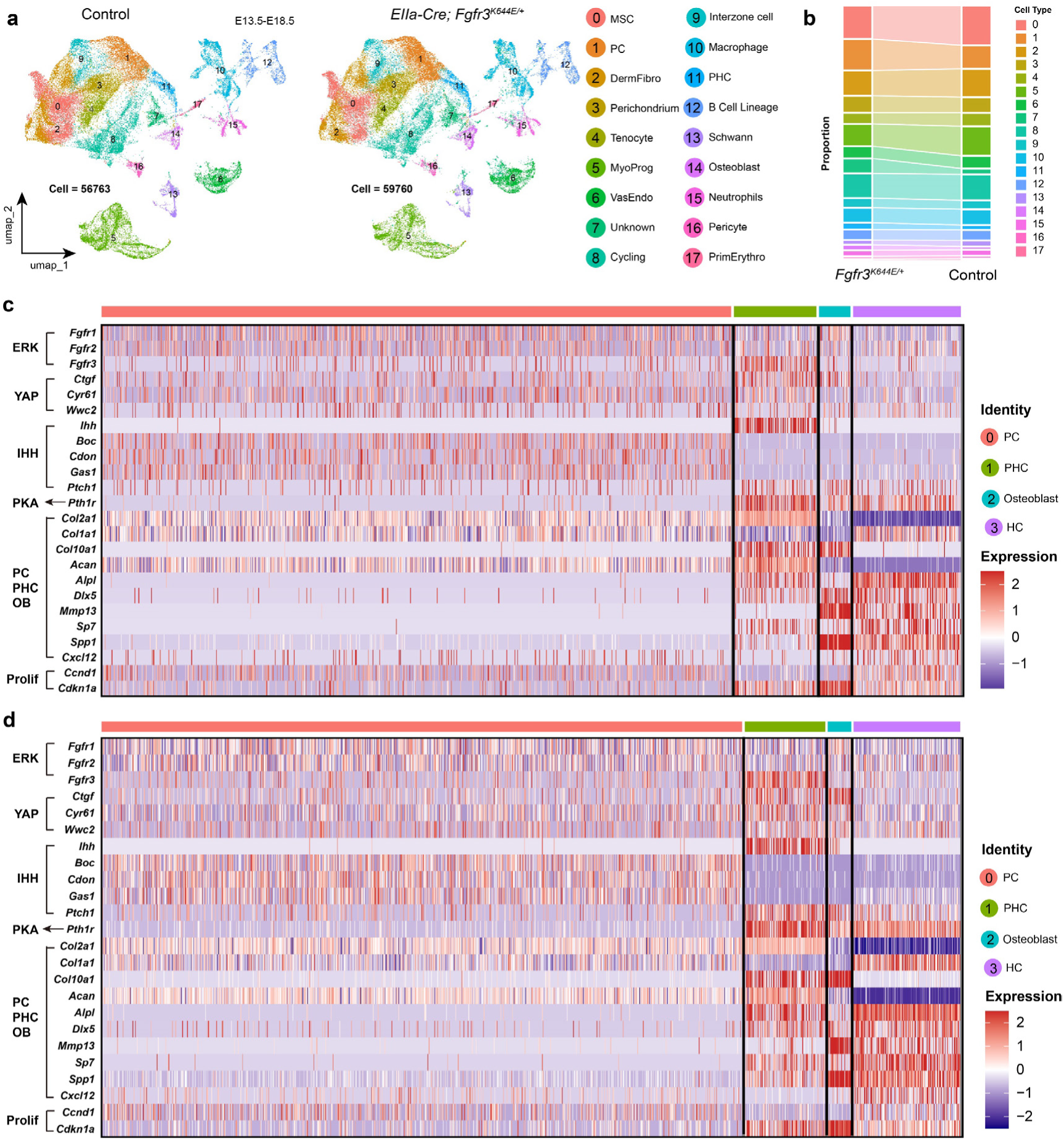
Gene express patterns in PCs, PHCs, HCs, and osteoblasts from E15.5 and E17.5 control mouse humeri. **a, b.** scRNA-seq of developing humeri from control and *Fgfr3-K644E* mice across six time points (E13.5– E18.5); n = 12 samples. MSCs, mesenchymal stem cell; DermFibro: Dermal fibroblast; MyoProg: Myogenic progenitor; VasEndo: Vascular endothelial cell; PrimErythro: Primitive erythroblast. **c, d.** Heatmap showing differentially expressed genes among PCs (including round and flattened PCs), PHCs, HCs, and osteoblasts (OBs) in control E15.5 (C) and E17.5 (D) humeri. Note that *Fgfr1* and *Fgfr2* were highly expressed in PCs but downregulated in PHCs, whereas *Fgfr3* expression peaked in PHCs. YAP signaling genes—*Cyr61* (*Ccn1*), *Ctgf* (*Ccn2*), and *Wwc2*— were also highly expressed in PHCs. In contrast, IHH coreceptors (Boc, Cdon, Gas1) were nearly exclusively expressed in PCs. Note that *Ccnd1* (a proliferation promoter) was decreased, whereas the differentiation factor *Cdkn1a* (*p21*) was increased in PHCs and HCs.

**Extended Data Fig. 9.**
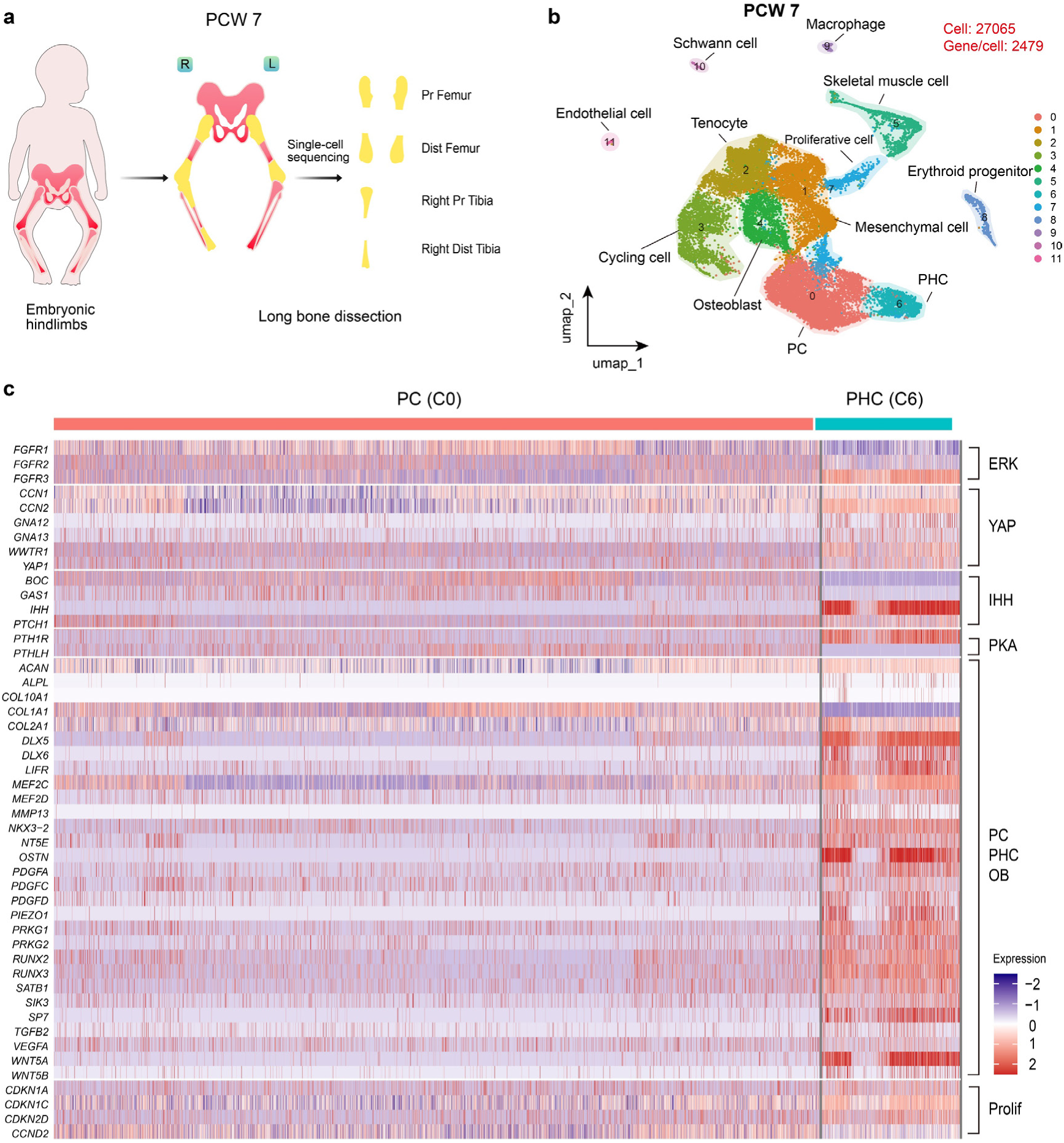
Gene expression patterns in PCs and PHCs from human GW9 femur and tibia. **a, b.** Reanalysis of a published scRNA-seq dataset: developing femur and tibia (proximal/distal) in a human fetus at GW9 (PCW7). **c.** Heatmap showing differentially expressed genes between PCs (including round and flattened PCs) and PHCs. Note that *FGFR1* and *FGFR2* were highly expressed in PCs but downregulated in PHCs, whereas *FGFR3* expression peaked in PHCs. YAP signaling genes—*CCN1* (*CYR61*), *CCN2* (*CTGF*), *GNA12/13*, *WWTR1* and *YAP1*— were also highly expressed in PHCs. IHH coreceptors (*BOC* and *GAS1*) were expressed almost exclusively in PCs, whereas both IHH and its repressor *PTCH1* were highly expressed in PHCs. *PTH1R* was expressed at lower levels in PCs but highly expressed in PHCs, whereas *PTHLH* was expressed in PCs, but not in PHC. The cell-cycle inhibitors *CDKN1A (p21)*, *CDKN1C* (*p57*), and *CDKN2D* (*p19*) were highly expressed in PHCs, consistent with their non-proliferative state.

**Extended Data Fig. 10.**
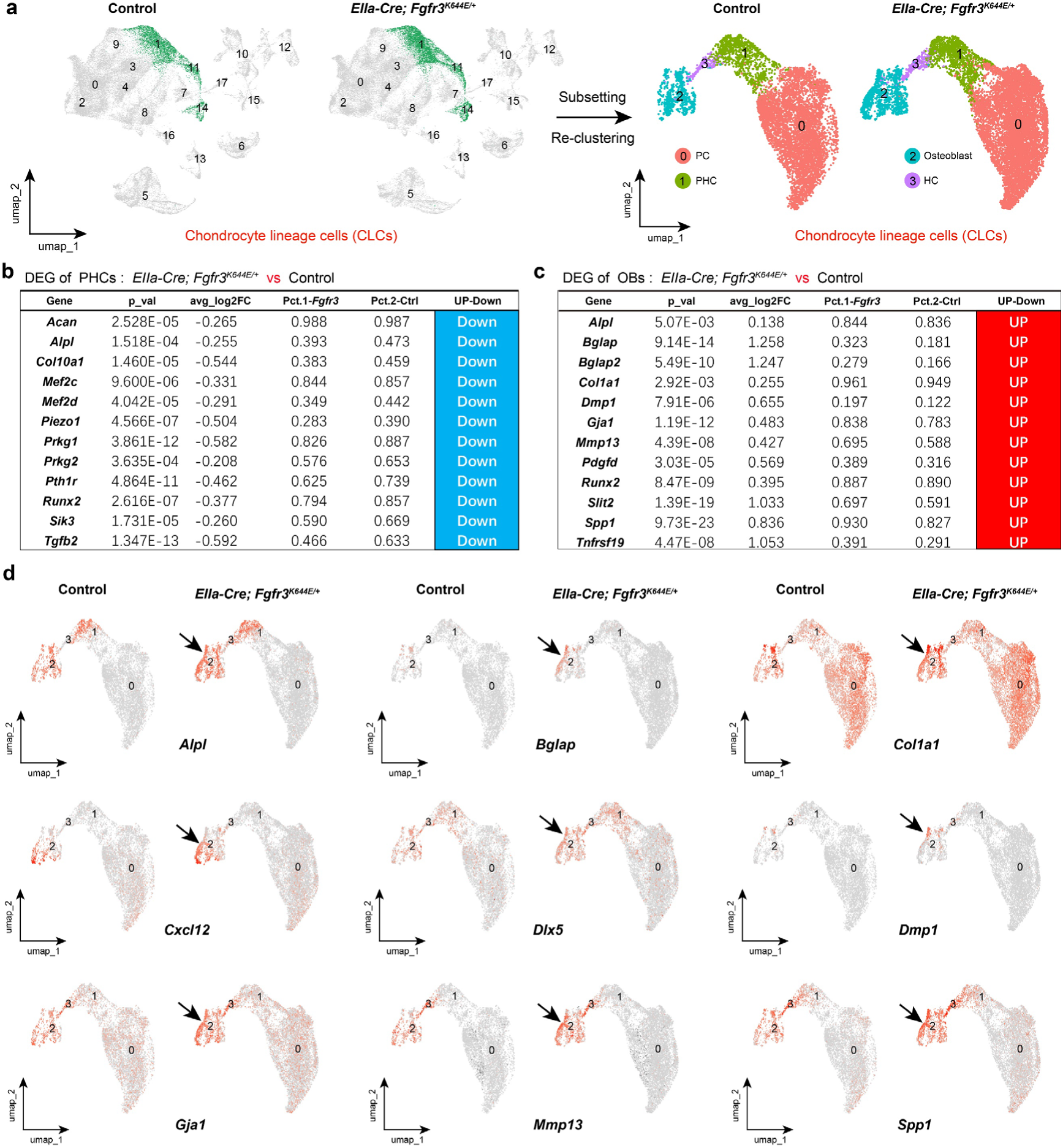
*Fgfr3-K644E* PHCs show immature gene expression, whereas osteoblasts appear precociously differentiated. **a.** A single-cell transcriptomic atlas of humerus development from E13.5 to E18.5 in control and *Fgfr3-K644E* mice was generated, followed by focused analysis of the chondrocyte lineage (PC, PHC, and HCs) and osteoblast populations. **b.** scRNA-Seq analysis revealed that *Fgfr3-K644E* PHC exhibit an immature state, characterized by downregulation of PHC-enriched genes. **c.** Osteoblasts (OBs) in *Fgfr3*-K*644*E mice underwent premature differentiation, marked by upregulation of late-stage markers (e.g., *Alpl*, *Bglap*, *Col1a1*). **d.** UMAP plots indicated the upregulation of genes associated with premature differentiation in osteoblasts (cluster 2) in *Fgfr3-K644E* mice (arrows).

**Extended Data Fig. 11.**
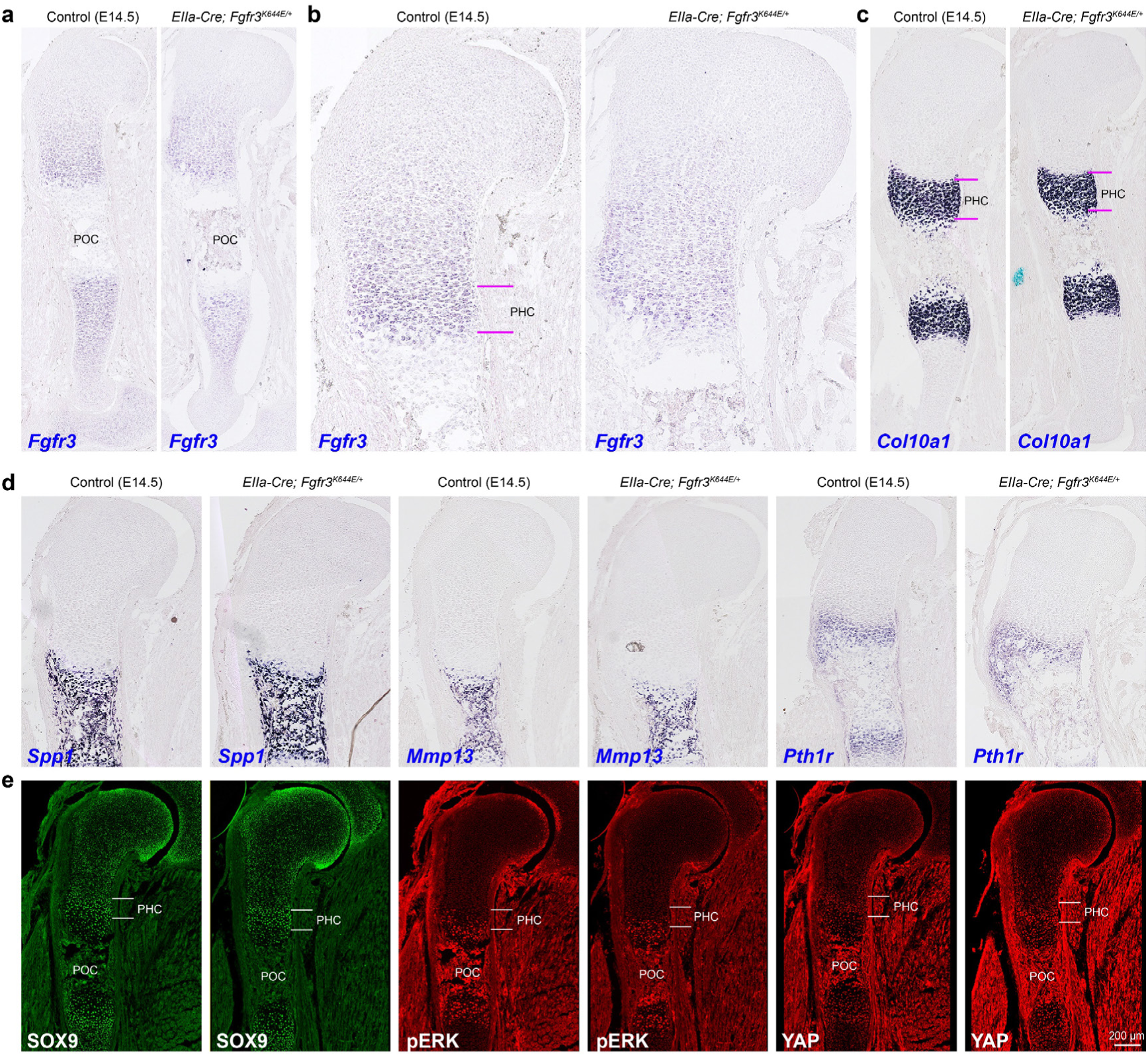
Humerus skeletal development showed only a mild phenotypic difference between *Fgfr3-K644E* and wild-type (WT) control mice at E14.5. **a-d.** The expression of *Fgfr3, Col10a1, Spp1, Mmp13*, and *Pth1R* in E14.5 humeri was examined by *in situ* hybridization. **e.** Representative immunostaining images showing expression patterns of SOX9, pERK, and YAP (detecting both YAP1 and TAZ antigens). Compared to wild-type (WT) controls, the *Fgfr3-K644E* humerus at E14.5 was slightly shorter (**c**).

**Extended Data Fig. 12.**
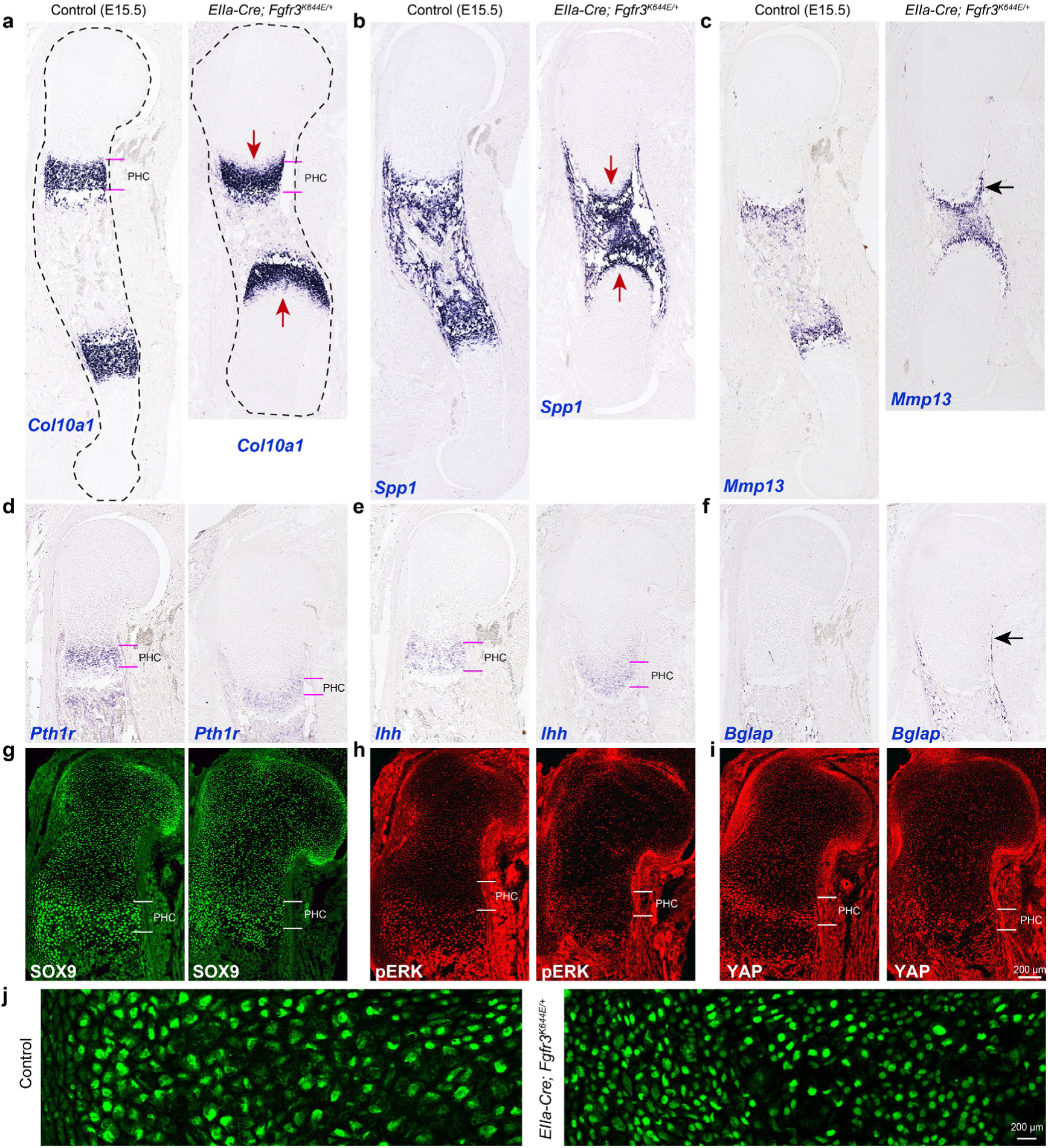
At E15.5, the development, differentiation, and maturation of PHCs in the humerus were impaired in *Fgfr3-K644E* mice relative to WT controls. **a-f.** *In situ* hybridization of *Col10a1*, *Spp1*, *Mmp13*, *Pth1r*, *Ihh,* and *Bglap* in E15.5 humeri revealed a marked delay in PHC maturation in *FGFR3-K644E* mice. Note that *Col10a1* expression in PHC was delayed (arrows in **a**), accompanied by a significantly reduced *Spp1* expression domain (arrows in **b**). *Pth1r* expression was significantly downregulated (**d**). These findings indicate that FGFR3-K644E suppresses PHC differentiation and maturation at E15.5. A mild increase in *Mmp13* and *Bglap* expression was observed in osteoblasts (arrows in **c, f**), indicating that the differentiation of HCs/osteoblasts was already occurring prematurely. These alterations contributed to a significantly shortened humerus in E15.5 *FGFR3-K644E* mice. **g-i.** Immunostaining of SOX9, pERK, and YAP. **j.** High-magnification imaging of SOX9+ cells showed enlarging PHCs in controls, whereas mutant PHCs remained undifferentiated with small, irregular nuclei.

**Extended Data Fig. 13.**
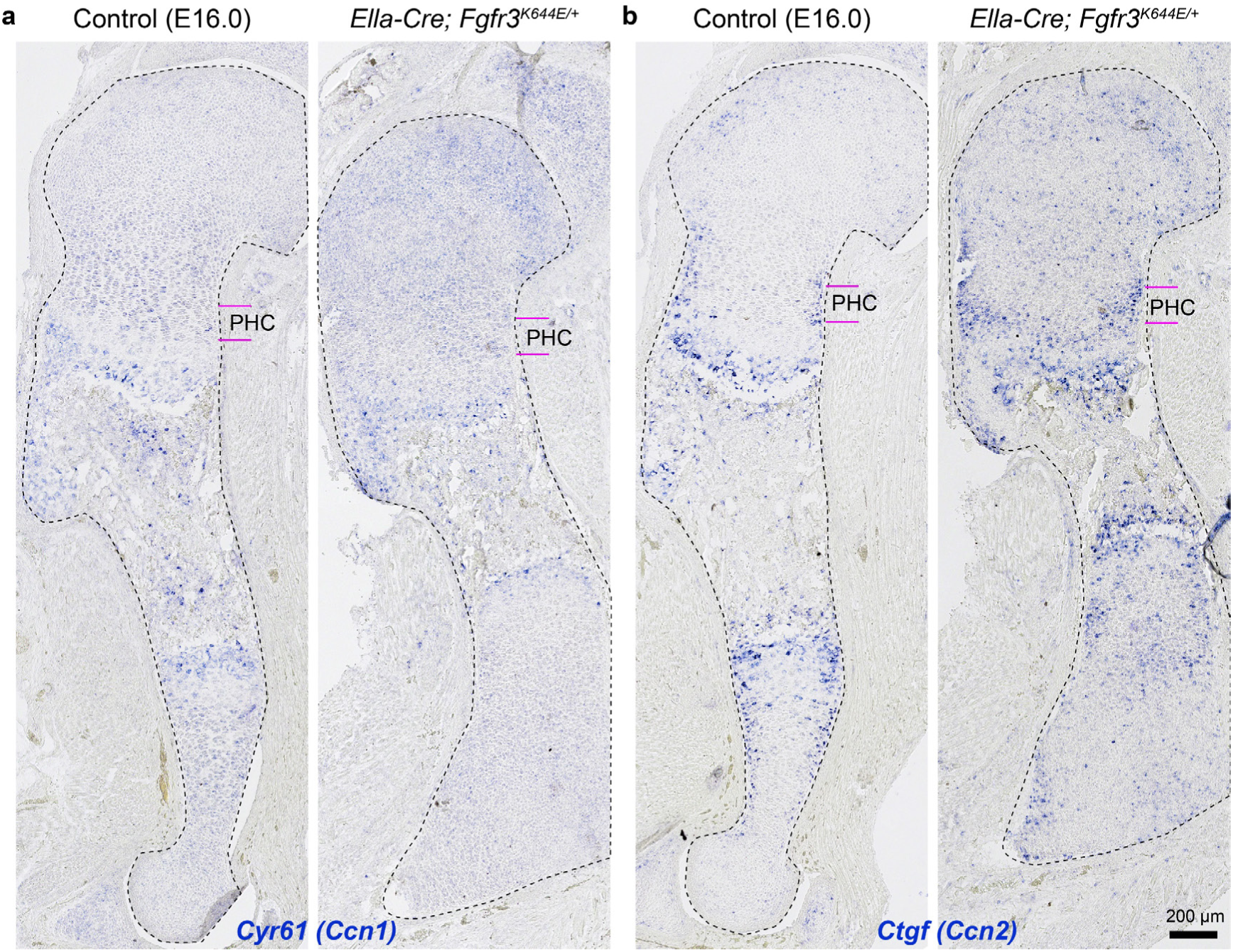
YAP signaling (YAP/TAZ activity) was examined in the mouse humerus at approximately E16.0. **a-g.** Analysis of *Cyr61* (*Ccn1*) and *Ctgf* (*Ccn2*) expression by *in situ* hybridization in E16.0 humerus sections. Compared with controls, *Cyr61* and *Ctgf* expression showed a slight but visible increase in *Fgfr3-K644E* mutant PHCs. Notably, despite this upregulation, *Cyr61*+ mutant chondrocytes maintained a small and irregular cell shape with disorganized morphology.

**Extended Data Fig. 14.**
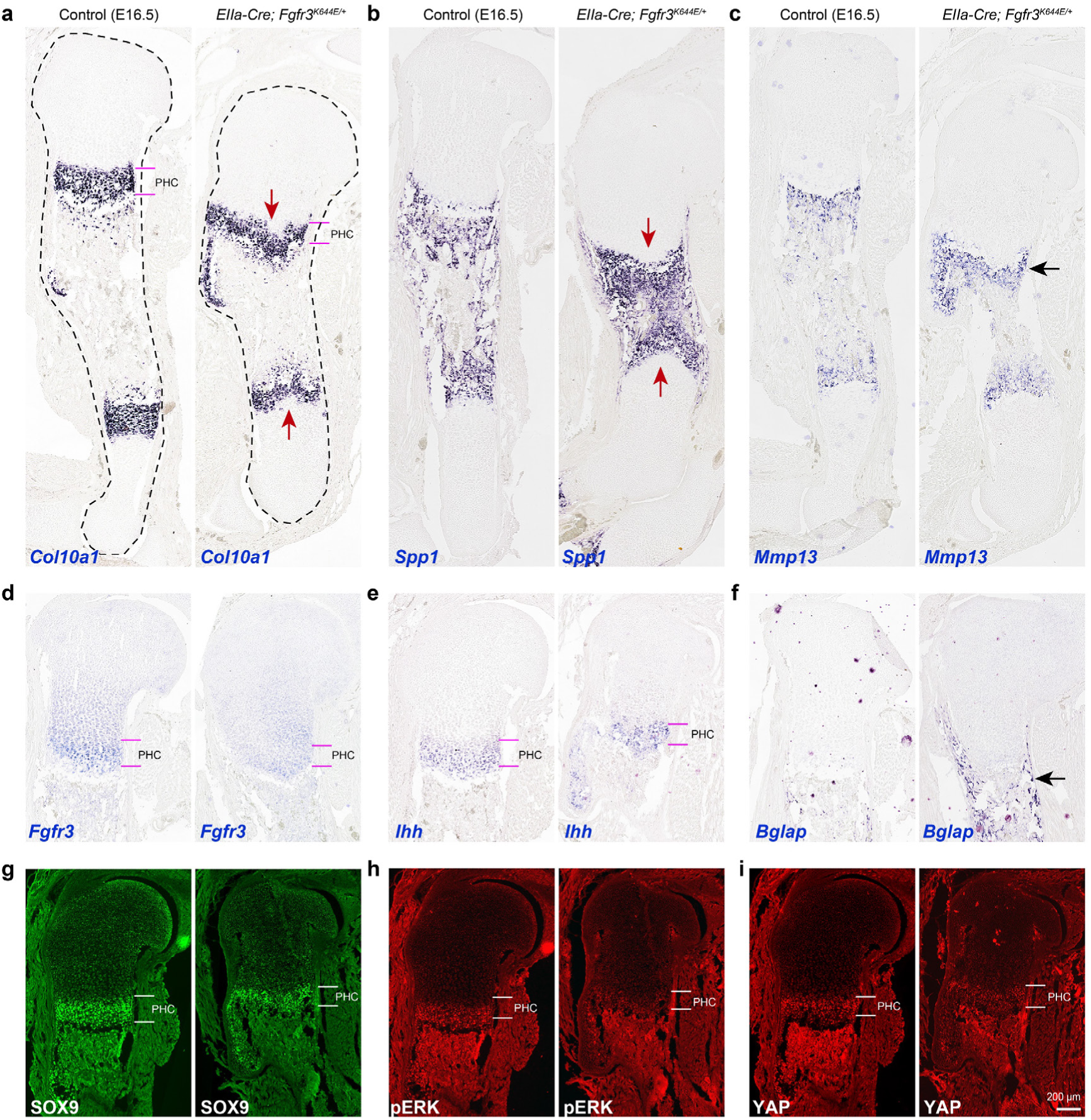
Humerus skeletal development was shortened and exhibited dwarfism in *Fgfr3*-*K644E* mice compared to wild-type (WT) controls at E16.5. **a-f.** *In situ* hybridization of E16.5 humeri revealed upregulation of *Spp1, Mmp13,* and *Bglap* in mutants (arrows). **g-i**. Immunofluorescence staining revealed a slight downregulation in SOX9, pERK and YAP protein levels in mutant PHCs/HCs compared with controls.

**Extended Data Fig. 15.**
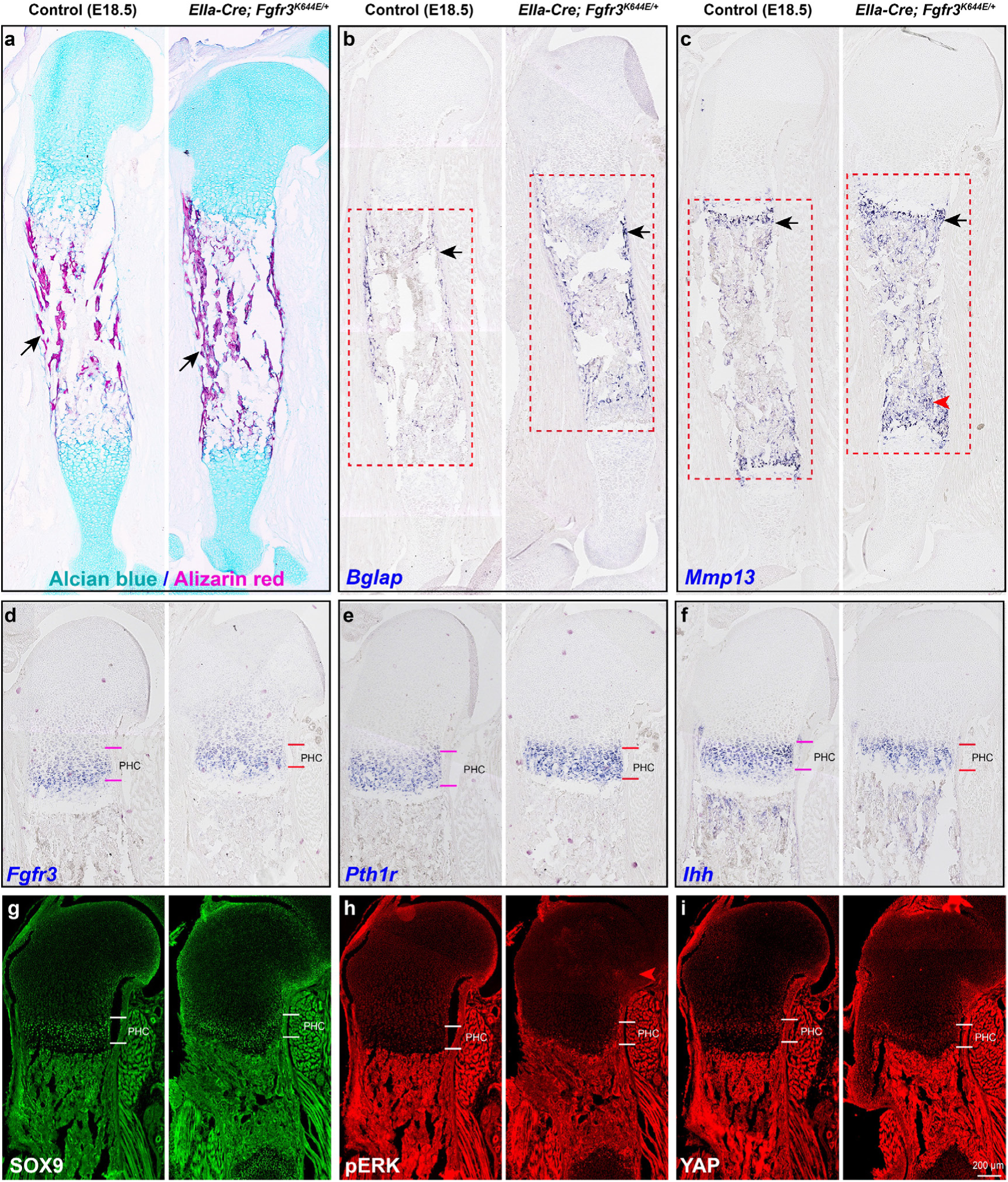
Osteoblasts in E18.5 *Fgfr3-K644E* mutant mice exhibited premature differentiation, accompanied by accelerated ossification. **a.** Alcian Blue (cartilage) / Alizarin Red (bone) staining of E18.5 humerus sections revealed premature ossification (arrows). **b-f.** *In situ* hybridization of E18.5 humeri revealed upregulation of *Bglap* and *Mmp13* in mutants (arrows in **b, c**). No significant differences were observed the expression of *Fgfr3*, *Pth1r*, and *Ihh*. **g-i**. Immunofluorescence staining indicated a reduction in SOX9+ and YAP+ PHCs in mutants relative to controls. In contrast, the pERK level did not differ significantly at E18.5.

**Table S1.** Antibodies used in this study.

| primary Ab | Species | Dilution | Manufacturer | Cat. No. |
| --- | --- | --- | --- | --- |
| HOPX | Rat | 1:800 | Oasis Biofarm | OB-PRT015 |
| HOPX | Rabbit | 1:800 | Proteintech | 11419-1-AP |
| NR2F1 | Mouse | 1:500 | PPMX | PP-H8132-00 |
| FOXJ1 | Mouse | 1:1000 | Invitrogen | 14-9965-80 |
| GFP | Chicken | 1:3000 | Aves labs | GFP-1020 |
| pERK1/2 | Rabbit | 1:400 | Cell Signaling | #4370 |
| SOX9 | Goat | 1:1000 | R&D Systems | AF3075 |
| SOX9 | Rabbit | 1:1000 | Abcam | ab185966 |
| YAP/TAZ | Rabbit | 1:1500 | Cell Signaling | 8418S |

**Table S2.** Single-cell RNA sequencing (scRNA-Seq) was performed on 12 mouse forelimb (humerus) samples and 8 cortical samples newly generated for this study.

| Number | Age | Genotype | Notes |
| --- | --- | --- | --- |
| 1 | E13.5 | *Fgfr3-K644E* mice | humerus |
| 2 | E13.5 | *Wild-type littermate* | humerus |
| 3 | E14.5 | *Fgfr3-K644E* mice | humerus |
| 4 | E14.5 | *Wild-type littermate* | humerus |
| 5 | E15.5 | *Fgfr3-K644E* mice | humerus |
| 6 | E15.5 | *Wild-type littermate* | humerus |
| 7 | E16.5 | *Fgfr3-K644E* mice | humerus |
| 8 | E16.5 | *Wild-type littermate* | humerus |
| 9 | E17.5 | *Fgfr3-K644E* mice | humerus |
| 10 | E17.5 | *Wild-type littermate* | humerus |
| 11 | E18.5 | *Fgfr3-K644E* mice | humerus |
| 12 | E18.5 | *Wild-type littermate* | humerus |
| 13 | E12.5 | *Fgfr3-K644E* mice | cortex |
| 14 | E12.5 | *Wild-type littermate* | cortex |
| 15 | E15.5 | *Fgfr3-K644E* mice | cortex |
| 16 | E15.5 | *Wild-type littermate* | cortex |
| 17 | E18.5 | *Fgfr3-K644E* mice | cortex |
| 18 | E18.5 | *Wild-type littermate* | cortex |
| 19 | P2 | *Emx1-Cre; Fgfr3<sup>K644E/K644E</sup>* | cortex |
| 20 | P2 | *Control littermate* | cortex |

**Table S3.**
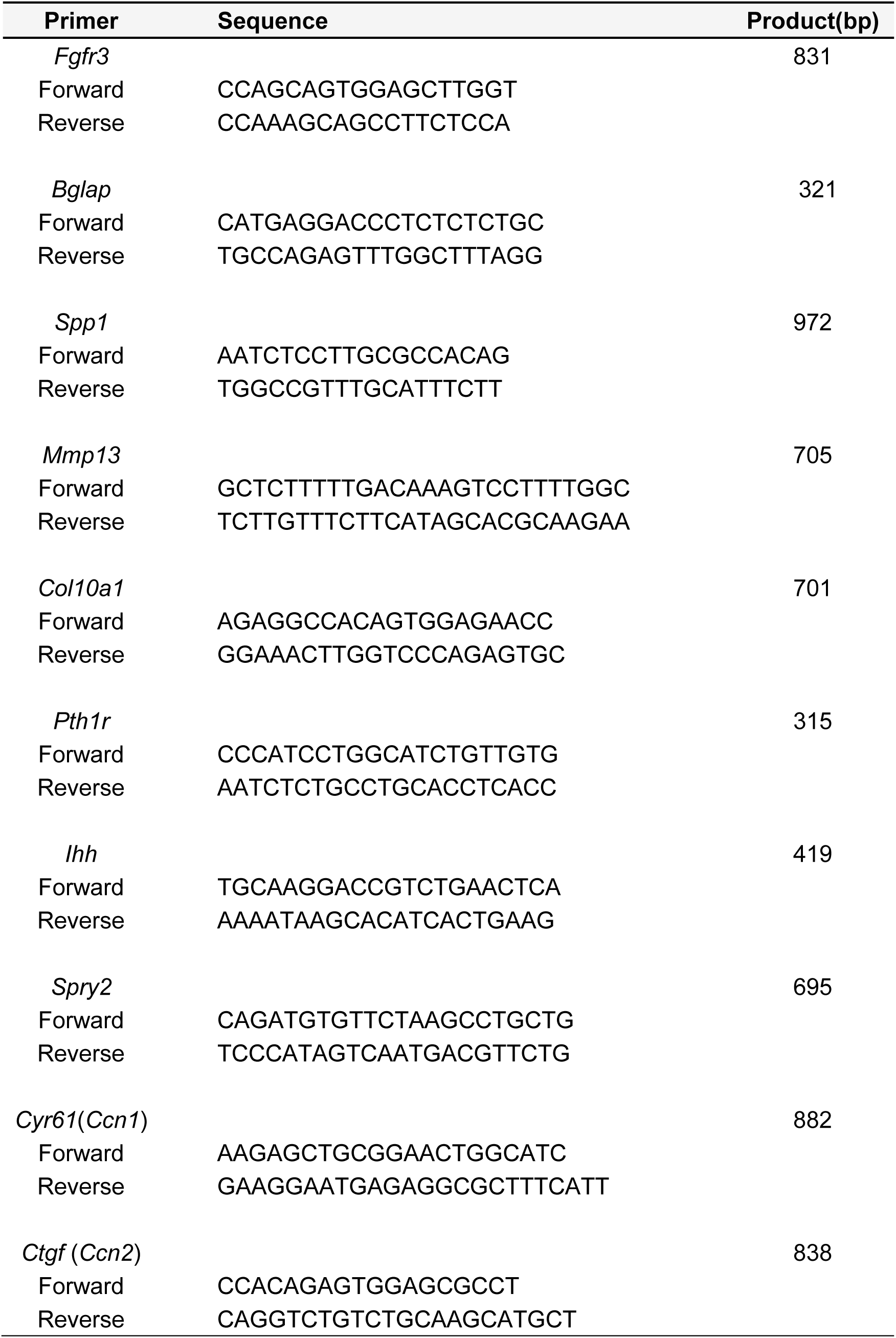
Primer pairs used in PCR preparing for in situ hybridization probes.

## References

1 Shiang, R. et al. Mutations in the transmembrane domain of FGFR3 cause the most common genetic form of dwarfism, achondroplasia. Cell 78, 335–342, doi:10.1016/0092-8674(94)90302-6 (1994).

2 Rousseau, F. et al. Mutations in the gene encoding fibroblast growth factor receptor-3 in achondroplasia. Nature 371, 252–254, doi:10.1038/371252a0 (1994).

3 Ornitz, D. M. & Legeai-Mallet, L. Achondroplasia: Development, pathogenesis, and therapy. Dev Dyn 246, 291–309, doi:10.1002/dvdy.24479 (2017).

4 Foldynova-Trantirkova, S., Wilcox, W. R. & Krejci, P. Sixteen years and counting: the current understanding of fibroblast growth factor receptor 3 (FGFR3) signaling in skeletal dysplasias. Hum Mutat 33, 29–41, doi:10.1002/humu.21636 (2012).

5 Kozhemyakina, E., Lassar, A. B. & Zelzer, E. A pathway to bone: signaling molecules and transcription factors involved in chondrocyte development and maturation. Development 142, 817–831, doi:10.1242/dev.105536 (2015).

6 Hevner, R. F. The cerebral cortex malformation in thanatophoric dysplasia: neuropathology and pathogenesis. Acta Neuropathol 110, 208–221, doi:10.1007/s00401-005-1059-8 (2005).

7 Tavormina, P. L. et al. Thanatophoric dysplasia (types I and II) caused by distinct mutations in fibroblast growth factor receptor 3. Nat Genet 9, 321–328, doi:10.1038/ng0395-321 (1995).

8 Iwata, T. & Hevner, R. F. Fibroblast growth factor signaling in development of the cerebral cortex. Dev Growth Differ 51, 299–323, doi:10.1111/j.1440-169X.2009.01104.x (2009).

9 Ornitz, D. M. & Itoh, N. The Fibroblast Growth Factor signaling pathway. Wiley Interdiscip Rev Dev Biol 4, 215–266, doi:10.1002/wdev.176 (2015).

10 Thomson, R. E., Pellicano, F. & Iwata, T. Fibroblast growth factor receptor 3 kinase domain mutation increases cortical progenitor proliferation via mitogen-activated protein kinase activation. J Neurochem 100, 1565–1578, doi:10.1111/j.1471-4159.2006.04285.x (2007).

11 Inglis-Broadgate, S. L. et al. FGFR3 regulates brain size by controlling progenitor cell proliferation and apoptosis during embryonic development. Developmental biology 279, 73–85, doi:10.1016/j.ydbio.2004.11.035 (2005).

12 Iwata, T. et al. A neonatal lethal mutation in FGFR3 uncouples proliferation and differentiation of growth plate chondrocytes in embryos. Hum Mol Genet 9, 1603–1613, doi:10.1093/hmg/9.11.1603 (2000).

13 Lin, T. et al. A central nervous system specific mouse model for thanatophoric dysplasia type II. Hum Mol Genet 12, 2863–2871, doi:10.1093/hmg/ddg309 (2003).

14 Sun, M. et al. ERK signaling expands mammalian cortical radial glial cells and extends the neurogenic period. Proc Natl Acad Sci U S A 121, e2314802121, doi:10.1073/pnas.2314802121 (2024).

15 Caronia-Brown, G., Yoshida, M., Gulden, F., Assimacopoulos, S. & Grove, E. A. The cortical hem regulates the size and patterning of neocortex. Development 141, 2855–2865, doi:10.1242/dev.106914 (2014).

16 Stork, P. J. & Schmitt, J. M. Crosstalk between cAMP and MAP kinase signaling in the regulation of cell proliferation. Trends Cell Biol 12, 258–266, doi:10.1016/s0962-8924(02)02294-8 (2002).

17 Cheloha, R. W., Gellman, S. H., Vilardaga, J. P. & Gardella, T. J. PTH receptor-1 signalling-mechanistic insights and therapeutic prospects. Nat Rev Endocrinol 11, 712–724, doi:10.1038/nrendo.2015.139 (2015).

18 Zhang, Z. et al. Integrated ERK-PKA-YAP/TAZ-SHH Signaling Orchestrates Cortical Radial Glia Identity and Lineage Diversification. Adv Sci (Weinh) 13, e13571, doi:10.1002/advs.202513571 (2026).

19 Hoffmann, R., Baillie, G. S., MacKenzie, S. J., Yarwood, S. J. & Houslay, M. D. The MAP kinase ERK2 inhibits the cyclic AMP-specific phosphodiesterase HSPDE4D3 by phosphorylating it at Ser579. Embo J 18, 893–903, doi:10.1093/emboj/18.4.893 (1999).

20 Lavado, A. et al. YAP/TAZ maintain the proliferative capacity and structural organization of radial glial cells during brain development. Developmental biology 480, 39–49, doi:10.1016/j.ydbio.2021.08.010 (2021).

21 Park, R. et al. Yap is required for ependymal integrity and is suppressed in LPA-induced hydrocephalus. Nat Commun 7, 10329, doi:10.1038/ncomms10329 (2016).

22 Lahti, L. et al. Sox9 and nuclear factor I transcription factors regulate the timing of neurogenesis and ependymal maturation in dopamine progenitors. Development 152, doi:10.1242/dev.204421 (2025).

23 Hoppner, J. et al. A mouse model of Jansen’s metaphyseal chondrodysplasia for investigating disease mechanisms and candidate therapeutics. Proc Natl Acad Sci U S A 122, e2500176122, doi:10.1073/pnas.2500176122 (2025).

24 Schipani, E., Kruse, K. & Juppner, H. A constitutively active mutant PTH-PTHrP receptor in Jansen-type metaphyseal chondrodysplasia. Science 268, 98–100, doi:10.1126/science.7701349 (1995).

25 Vortkamp, A. et al. Regulation of rate of cartilage differentiation by Indian hedgehog and PTH-related protein. Science 273, 613–622, doi:10.1126/science.273.5275.613 (1996).

26 Kong, J. H., Siebold, C. & Rohatgi, R. Biochemical mechanisms of vertebrate hedgehog signaling. Development 146, doi:10.1242/dev.166892 (2019).

27 Hui, C. C. & Angers, S. Gli proteins in development and disease. Annu Rev Cell Dev Biol 27, 513–537, doi:10.1146/annurev-cellbio-092910-154048 (2011).

28 Rao, R., Salloum, R., Xin, M. & Lu, Q. R. The G protein Galphas acts as a tumor suppressor in sonic hedgehog signaling-driven tumorigenesis. Cell Cycle 15, 1325–1330, doi:10.1080/15384101.2016.1164371 (2016).

29 Yu, F. X. et al. Protein kinase A activates the Hippo pathway to modulate cell proliferation and differentiation. Genes Dev 27, 1223–1232, doi:10.1101/gad.219402.113 (2013).

30 Lawrence, J. E. G. et al. Single-cell transcriptomics identifies chondrocyte differentiation dynamics in vivo and in vitro. Dev Cell 60, 3066–3084 e3068, doi:10.1016/j.devcel.2025.06.031 (2025).

31 Hong, J. H. et al. TAZ, a transcriptional modulator of mesenchymal stem cell differentiation. Science 309, 1074–1078, doi:10.1126/science.1110955 (2005).

32 Zarka, M., Hay, E. & Cohen-Solal, M. YAP/TAZ in Bone and Cartilage Biology. Front Cell Dev Biol 9, 788773, doi:10.3389/fcell.2021.788773 (2021).

33 Varelas, X. et al. The Crumbs complex couples cell density sensing to Hippo-dependent control of the TGF-beta-SMAD pathway. Dev Cell 19, 831–844, doi:10.1016/j.devcel.2010.11.012 (2010).

34 Faedo, A., Borello, U. & Rubenstein, J. L. Repression of Fgf signaling by sprouty1-2 regulates cortical patterning in two distinct regions and times. J Neurosci 30, 4015–4023, doi:10.1523/JNEUROSCI.0307-10.2010 (2010).

35 Garel, S., Huffman, K. J. & Rubenstein, J. L. Molecular regionalization of the neocortex is disrupted in Fgf8 hypomorphic mutants. Development 130, 1903–1914, doi:10.1242/dev.00416 (2003).

36 Fukuchi-Shimogori, T. & Grove, E. A. Emx2 patterns the neocortex by regulating FGF positional signaling. Nat Neurosci 6, 825–831, doi:10.1038/nn1093 (2003).

37 Coletti, A. M. et al. Characterization of the ventricular-subventricular stem cell niche during human brain development. Development 145, doi:10.1242/dev.170100 (2018).

38 Spassky, N. et al. Adult ependymal cells are postmitotic and are derived from radial glial cells during embryogenesis. J Neurosci 25, 10–18 (2005).

39 St-Jacques, B., Hammerschmidt, M. & McMahon, A. P. Indian hedgehog signaling regulates proliferation and differentiation of chondrocytes and is essential for bone formation. Genes Dev 13, 2072–2086, doi:10.1101/gad.13.16.2072 (1999).

40 Gao, B. & Yang, Y. Planar cell polarity in vertebrate limb morphogenesis. Curr Opin Genet Dev 23, 438–444, doi:10.1016/j.gde.2013.05.003 (2013).

41 Vanyai, H. K. et al. Control of skeletal morphogenesis by the Hippo-YAP/TAZ pathway. Development 147, doi:10.1242/dev.187187 (2020).

42 Goto, H. et al. Loss of Mob1a/b in mice results in chondrodysplasia due to YAP1/TAZ-TEAD-dependent repression of SOX9. Development 145, doi:10.1242/dev.159244 (2018).

43 Hu, Y. J. et al. Piezo1-mediated mechanotransduction controls osteocyte maturation and dendrite development via a YAP-CCN-Src signaling axis. Nat Commun 16, 10859, doi:10.1038/s41467-025-65636-9 (2025).

44 Cooper, K. L. et al. Multiple phases of chondrocyte enlargement underlie differences in skeletal proportions. Nature 495, 375–378, doi:10.1038/nature11940 (2013).

45 Regard, J. B. et al. Activation of Hedgehog signaling by loss of GNAS causes heterotopic ossification. Nat Med 19, 1505–1512, doi:10.1038/nm.3314 (2013).

46 Lakso, M. et al. Efficient in vivo manipulation of mouse genomic sequences at the zygote stage. Proc Natl Acad Sci U S A 93, 5860–5865, doi:10.1073/pnas.93.12.5860 (1996).

47 Gorski, J. A. et al. Cortical excitatory neurons and glia, but not GABAergic neurons, are produced in the Emx1-expressing lineage. J Neurosci 22, 6309–6314 (2002).

48 Tronche, F. et al. Disruption of the glucocorticoid receptor gene in the nervous system results in reduced anxiety. Nat Genet 23, 99–103, doi:10.1038/12703 (1999).

49 Srinivasan, L. et al. PI3 kinase signals BCR-dependent mature B cell survival. Cell 139, 573–586, doi:10.1016/j.cell.2009.08.041 (2009).

50 Ivkovic, S. et al. Connective tissue growth factor coordinates chondrogenesis and angiogenesis during skeletal development. Development 130, 2779–2791, doi:10.1242/dev.00505 (2003).

51 Rigueur, D. & Lyons, K. M. Whole-mount skeletal staining. Methods Mol Biol 1130, 113–121, doi:10.1007/978-1-62703-989-5_9 (2014).

52 Wallin, J. et al. The role of Pax-1 in axial skeleton development. Development 120, 1109–1121, doi:10.1242/dev.120.5.1109 (1994).

53 Telley, L. et al. Sequential transcriptional waves direct the differentiation of newborn neurons in the mouse neocortex. Science 351, 1443–1446, doi:10.1126/science.aad8361 (2016).

54 Govindan, S., Oberst, P. & Jabaudon, D. In vivo pulse labeling of isochronic cohorts of cells in the central nervous system using FlashTag. Nat Protoc 13, 2297–2311, doi:10.1038/s41596-018-0038-1 (2018).

55 Zhang, Q. et al. The Zinc Finger Transcription Factor Sp9 Is Required for the Development of Striatopallidal Projection Neurons. Cell Rep 16, 1431–1444, doi:10.1016/j.celrep.2016.06.090 (2016).

